# ASTN2 modulates synaptic strength by trafficking and degradation of surface proteins

**DOI:** 10.1101/337618

**Authors:** Hourinaz Behesti, Taylor Fore, Peter Wu, Zachi Horn, Mary Leppert, Court Hull, Mary E Hatten

## Abstract

Surface protein dynamics dictate synaptic connectivity and function in neuronal circuits. *ASTN2*, a gene disrupted by copy number variations (CNVs) in neurodevelopmental disorders, including autism spectrum, was previously shown to regulate the surface expression of ASTN1 in glial-guided neuronal migration. Here, we demonstrate that ASTN2 binds to and regulates the surface expression of multiple synaptic proteins in post-migratory neurons by endocytosis, resulting in modulation of synaptic activity. In cerebellar Purkinje cells (PCs), by immuno-gold electron microscopy, ASTN2 localizes primarily to endocytic and autophagocytic vesicles in the cell soma and in subsets of dendritic spines. Overexpression of ASTN2 in PCs, but not of ASTN2 lacking the FNIII-domain commonly disrupted by CNVs in patients including in a family presented here, increases inhibitory and excitatory postsynaptic activity and reduces levels of ASTN2 binding partners. Our data suggest a fundamental role for ASTN2 in dynamic regulation of surface proteins by endocytic trafficking and protein degradation.

## Introduction

ASTN2 is a large vertebrate-specific transmembrane protein, expressed in the developing and adult brain, with the highest levels detected in the cerebellum (1). Previously, we showed that ASTN2 interacts with ASTN1, a surface membrane protein that regulates glial-guided neuronal migration (1–4). Recently, copy number variations (CNVs) of *ASTN2*, both deletions and duplications (see Fig. S1), were identified in patients with neurodevelopmental disorders (NDDs) including autism spectrum disorder (ASD), schizophrenia, attention-deficit/hyperactivity disorder (ADHD), bipolar disease, intellectual disability (ID), and global developmental delay (5–9). In particular, *ASTN2* CNVs mainly affected the MAC/Perforin (MACPF) and the FNIII encoding regions of the gene and were identified as a significant risk factor for ASD in males in a study of 89,985 subjects (10).

Despite shared protein homology, ASTN2 but not ASTN1, is highly expressed in the adult cerebellum, long after completion of neuronal migration, suggestive of key additional roles unrelated to migration. While the cerebellum has traditionally been associated with motor control, recent evidence has suggested non-motor functions including language, visuospatial memory, attention, and emotion (11–13). In particular, loss of cerebellar Purkinje cells (PCs), is one of the most consistent findings in post-mortem studies in ASD patients (14). Moreover, specific targeting of cerebellar neurons in mouse models of ASD-associated genes, leads to impaired cerebellar learning (15) and social behaviors (16). The mechanism of action of ASTN2 in post-migratory neurons and how it may contribute to the pathophysiology of NDDs is currently unknown.

Here, we describe a family with a paternally inherited intra-genic *ASTN2* duplication and NDD including ASD, and most notably learning difficulty and speech and language delay. By immuno-gold electron microscopy (EM), we show that ASTN2 localizes primarily to vesicles in PC soma and to subsets of dendritic spines. By immunoprecipitation/mass spectrometry (IP/mass spec), we identify ASTN2 binding partners including C1q, Neuroligins, ROCK2, and SLC12a5 (KCC2), and show that ASTN2 removes surface proteins by endocytosis. Further, ASTN2 is found in a subset of vesicles along the entire endosomal pathway and links to the endosomal trafficking machinery via binding to the adaptor protein AP-2 and the vacuolar protein-sorting-associated protein 36 (VPS36). Importantly, consistent with a role in regulating the surface expression of key synaptic proteins, while conditional overexpression of ASTN2 in PCs increases synaptic strength, ASTN2 with deletion of the FNIII domain, the region commonly disrupted by CNVs in patients including the family presented here, is inefficient at changing synaptic activity. At the molecular level, overexpression of ASTN2 results in reduced protein levels of its synaptic binding partners. Our study identifies ASTN2 as a molecule that modulates the composition of the surface membrane proteome. We propose that the intra-genic *ASTN2* CNVs in patients result in misregulation of surface protein turnover, which is crucial for normal synaptic activity.

## Results

### Paternally inherited ASTN2 CNV in a family with ASD, ID, and speech and language delay

Single nucleotide polymorphism (SNP) array genetic testing of a child presented at 19 months of age, identified a 171 kb duplication at 9q33.1, affecting exons 16-19 of *ASTN2* (personal communication, L. Jamal, Johns Hopkins). The CNV was present in the father and 3/5 children, indicating a paternally inherited heterozygous duplication. The children displayed a range of NDDs (Table S1) including ID and ASD. Two features in particular stood out in the affected children, namely learning difficulty and speech and language delay regardless of other diagnoses.

To investigate how the duplication of exons 16-19, which code for part of the MACPF and the FNIII domains of ASTN2, affects ASTN2 expression, we obtained peripheral blood mononuclear cells (PBMCs) from patients, where *ASTN2* expression was detected in the CD4+ T cell fraction (Fig. 1a). The duplication was predicted to either result in an mRNA encoding an intact MACPF domain but a truncated FNIII domain due to the creation of a frameshift stop codon (termed JDUP, Fig. S2), or nonsense-mediated decay of the mRNA. In CD4+ T cells isolated from 2/3 of the boys with the *ASTN2* CNV as well as the father, *ASTN2* was reduced by ∼30-50% compared to controls including the mother (Fig. 1b). We detected two protein bands, one of which is absent in the mouse, both of which were ∼50% lower in patients compared to controls by Western blot (Fig. 1c). While the mRNA quantification (Fig. 1b) suggests that the majority of the duplicated mRNA undergoes nonsense-mediated decay, we cannot exclude that low levels of the truncated protein (termed JDUP) is expressed in patients, as the antibody used does not recognize JDUP (verified with a deletion construct, data not shown).

**Figure 1.**
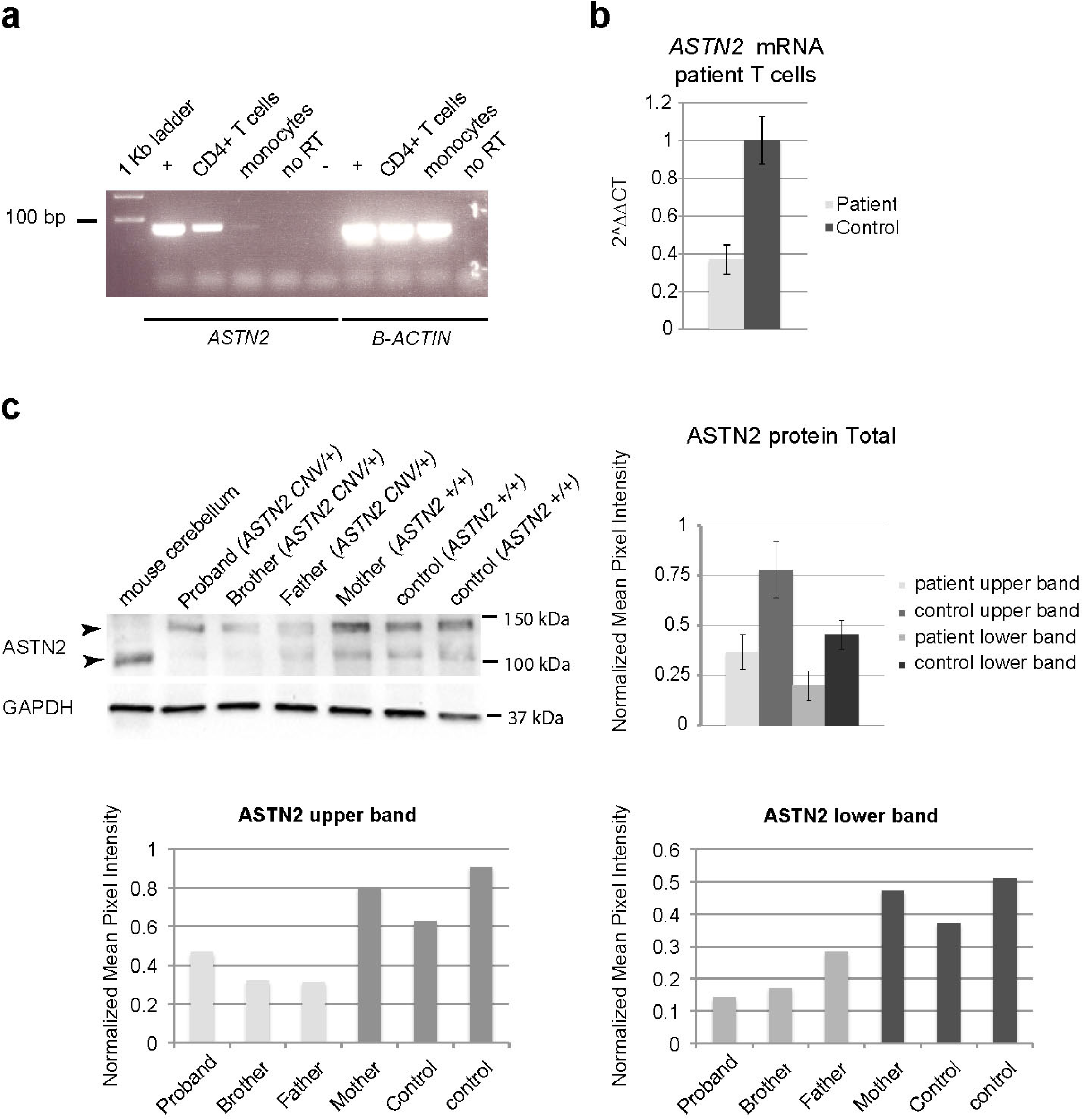
ASTN2 expression in patients. **(a)** Expression of *ASTN2* detected in human CD4+ T cells but not in monocytes. Positive (human fibroblasts), negative (no template), and no reverse transcription (no RT) controls are indicated. **b**) *ASTN2* mRNA levels, expressed as 2^^ΔΔCT^ (cycle time by qRT-PCR) in relation to GUSB (endogenous control) and **(c)** protein levels in *ASTN2* CNV patient T cells versus controls, quantified in graph to the right. Quantifications of individual ASTN2 bands (upper and lower bands) in relation to GAPDH are shown at the bottom. N= 3 patients and 3 controls. Bars show means +/−1 S.D. (standard deviation).

In the DECIPHER database, which currently contains clinical and genetic information on 11,887 NDD patients, ID is reported in 11/18 (61%) patients with *ASTN2* CNVs (both deletions and duplications); a slightly higher rate of occurrence than in the overall NDD population (6735/11887, 57%). Speech and language delay was reported in 5/18 *ASTN2* CNV patients (28%), also above the rate observed in the overall NDD population (2505/11887, 21%). In relation to other genes that are highly associated with either ID or ASD/ID, *ASTN2* CNV patients fell above the median for ID among the investigated genes (above 6/8 ASD-associated genes and within the range of the ID genes), and on the median among these genes for speech and language delay (Fig. S1b). Thus, ID is the single most commonly occurring feature in patients with *ASTN2* CNVs (both deletions and duplications) followed by speech and language delay, including in patients diagnosed with ASD.

### ASTN2 protein localization in the juvenile brain

To investigate the function of ASTN2, we first analysed its subcellular localization in the mouse cerebellum; the strongest site of expression in the brain (1). Immunohistochemistry (for antibody validation see Fig. S3a, b and (1)) in the juvenile mouse cerebellum showed ASTN2 in granule cells (GC), in the molecular layer, and at higher levels in PCs (Fig. 2a). In PCs, punctate labeling was detected in the PC body, in the dendritic stalk, and in dendrites (Fig. 2c-e). Immuno-EM labeling revealed that ASTN2 localized to membranes in the ER, and small round trafficking vesicles near the ER, the Golgi, and the plasma membrane of PCs (Fig. 2g). ASTN2 also localized to endocytic vesicles (Fig. 2g-i), and autophagosomes (Fig. 2g, j). A subset of dendritic spines, mostly in proximal regions of PCs, was positive for ASTN2. In labeled spines, ASTN2 localized to membranes near, but not directly at, the postsynaptic density (Fig. 2l-o). Co-labeling of ASTN2 with recycling (Rab4), early (Rab5), and late (Rab7) endosomal markers revealed that a small subset of ASTN2 puncta localize to all these fractions of the endocytic pathway in the juvenile cerebellum (Fig. S4). The punctate expression pattern of ASTN2 and its localization to membranes of endocytic and autophagocytic vesicles suggests involvement in trafficking and its presence proximal to synapses in post-migratory neurons raised the possibility that ASTN2 is involved in synaptic function.

**Figure 2.**
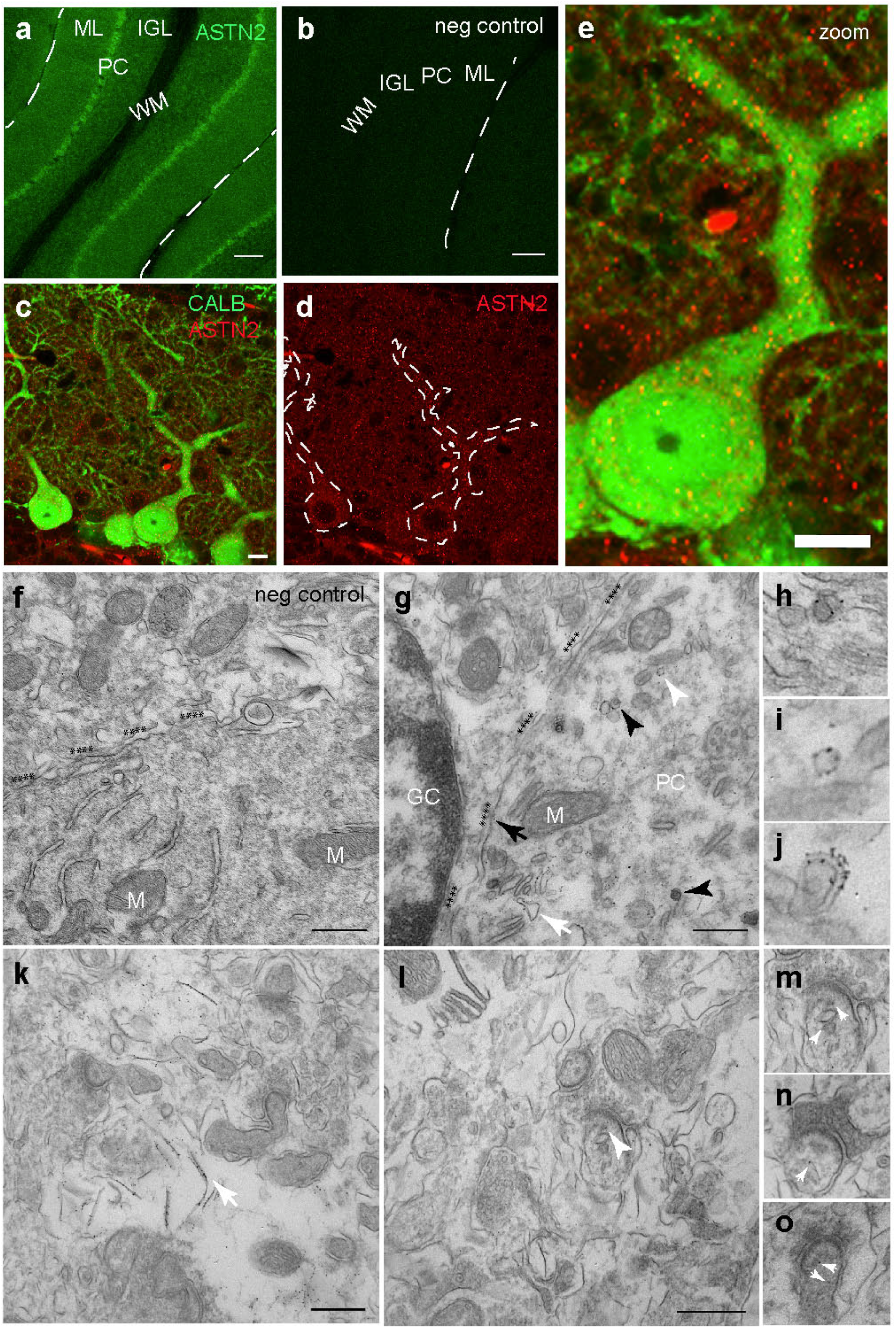
ASTN2 subcellular protein localization in the cerebellum. Sagittal sections of cerebellum labeled with antibodies against (**a**) ASTN2 (green) and (**c-e**) ASTN2 (red) and Calbindin (green) at P15. (b) negative control (no primary antibody), (e) zoom of one PC. Dotted lines in (d) outline PC bodies and primary dendrites. (**f-o**) Immuno-gold EM labeling of ASTN2 at P28. (f) Negative control (no primary), (g) ASTN2 labeling in a PC soma associated with the plasma membrane (highlighted by asterisks and black arrow), membranes of the ER (white arrow), trafficking vesicles (white arrowhead and, see also **i**), and autophagosomes (black arrowheads and see also **j**). (h) High power image showing ASTN2 labeling associated with an endocytic vesicle at the plasma membrane. (k) PC dendrite in the ML with ASTN2 labeling on isolation membranes (arrow). Synapses in this image are negative for ASTN2. (l) PC dendritic area with positive labeling in a spine (white arrowhead). (m-o) higher magnification examples of PC dendritic spines showing ASTN2 labeling (arrows). N= 3 biological and 3 technical replicates. GC, granule cell; IGL, internal granule layer; M, mitochondria; ML, molecular layer; PC, Purkinje cell; WM, white matter. Scale bars: 100 μm in a, b, 10 μm in c-e, 0.5 μm in f, g, k, l.

### ASTN2 binds to and reduces the surface expression of synaptic proteins by endocytosis

To investigate whether ASTN2 has a synaptic role, we first examined if it binds to key adhesion proteins known to regulate PC synaptic function. Co-immuno precipitation (IP) experiments revealed that ASTN2 interacts with members of the Neuroligin family, as does the truncated JDUP version (Fig. 3a). By Western blot, NLGN1/2 interacted more strongly with ASTN2 than with JDUP, while NLGN3/4 interacted more strongly with JDUP than with ASTN2. Thus ASTN2 binds to Neuroligins and the presence of the FNIII domain differentially impacts the affinity of ASTN2 for different binding partners.

**Figure 3.**
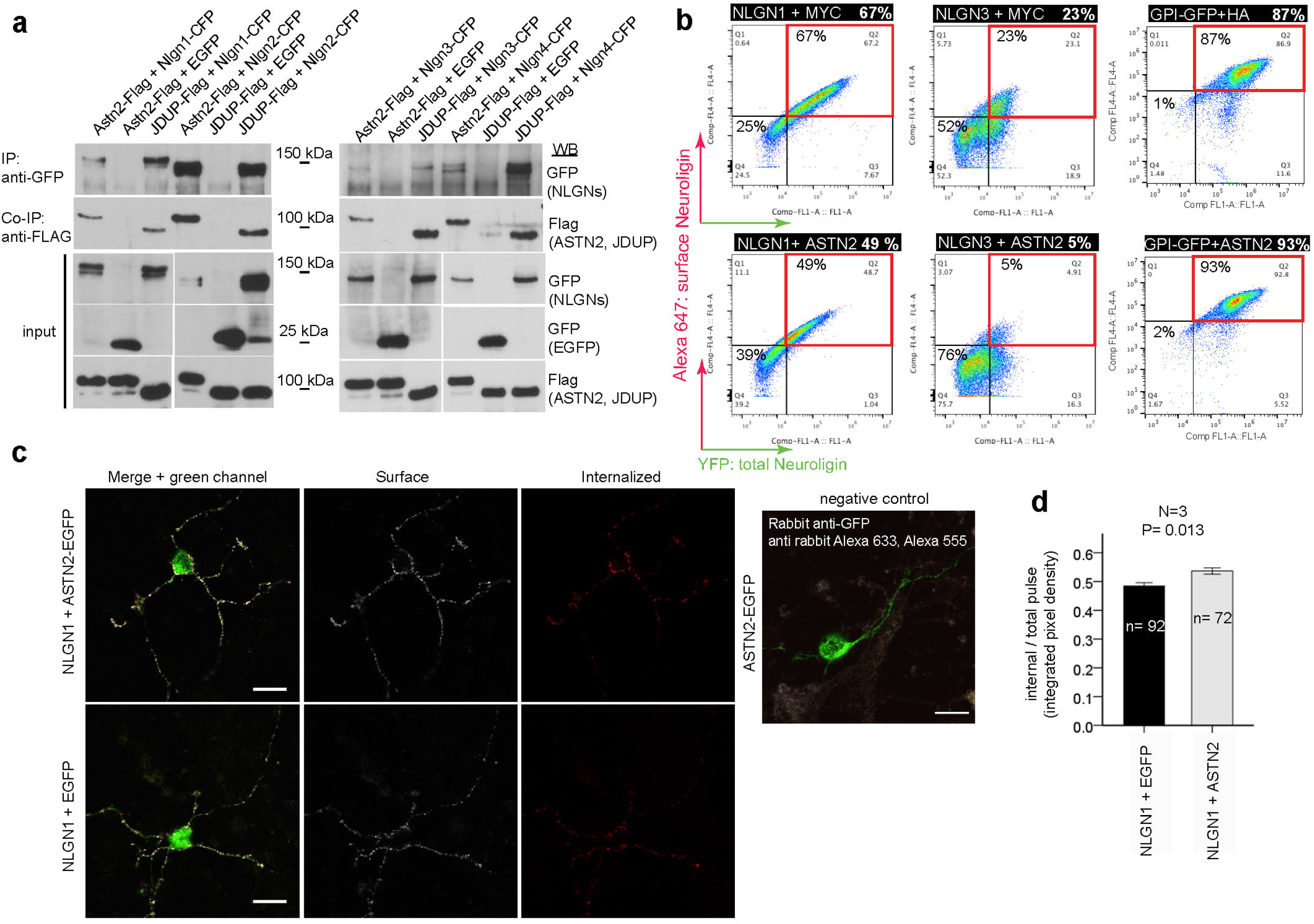
ASTN2 regulation of Neuroligin surface expression by protein-protein binding and endocytosis. **(a)** Western blots showing co-IP of ASTN2 and JDUP with Neuroligins 1-4 in HEK 293T. **(b)** Live immuno-labeling of surface Neuroligin expression (Alexa-647, red quadrants) in HEK 293T cells analysed by flow cytometry in cells co-expressing NLGN1-HA-YFP or NLGN3-YFP with either a MYC control vector (top panels) or with ASTN2-HA (bottom panels). Surface GPI-anchored EGFP is unaltered by ASTN2 (far right graphs). **(c)** Pulse-chase labeling of NLGN1-HA-YFP co-expressed with either EGFP or ASTN2-EGFP in GCs showing surface (white) and internalized (red) NLGN1 labeling after a 20 min chase. Far right image is a negative control, showing that the EGFP from ASTN2-EGFP (or EGFP) is not detected on the surface. **(d)** Quantification of the pulse-chase expressed as integrated pixel density (sum of all pixel intensities per area minus the background) of the internal labeling divided by the integrated pixel density of the total pulse (red + white). Graph show means +/−1 SEM. N= number of experiments, n= total number of cells analysed. P-value calculated by ANCOVA (see methods). Protein ladder in kDa. WB, Western blot, Scale bars: 10μm.

To investigate whether ASTN2 regulates the surface expression of Neuroligins, we quantified the surface expression of NLGN1-EGFP by live immunolabeling and flow cytometry in the presence and absence of ASTN2 and found a reduction in surface NLGN1 in cells co-transfected with ASTN2 as compared to cells without (49% versus 67%, Fig. 3b). This reduction was even more marked for NLGN3, in the presence of ASTN2 (Fig. 3b, 23% versus 5%). However, ASTN2 did not reduce the surface expression of glycosylphosphatidylinositol (GPI)-anchored surface EGFP (Fig. 3b). Hence ASTN2 specifically removes Neuroligins from the surface of HEK cells due to protein-protein interaction.

We then examined whether ASTN2 expression reduced surface NLGN1 also in neurons. For these experiments we chose to target GCs, which are far more numerous than PCs and also express ASTN2, and together with PCs, are the main neuronal subtype in the cerebellum. We used overexpression to disrupt the stoichiometry of ASTN2 protein complexes, as knockdown of ASTN2 protein was not possible in neurons (Fig. S3), possibly due to the extremely long half-life of ASTNs in the brain (17). As seen in HEK293T cells, GCs grown in culture for 14 days and co-transfected with *Nlgn1* and *Astn2* had reduced surface expression of NLGN1 compared to controls by live immunolabeling of surface NLGN1-EGFP (Fig. S5a, b). To investigate whether the reduction in surface expression is due to endocytosis versus potential changes in surface insertion of NLGN1 upon expression from plasmids, we carried out pulse-chase labeling of surface NLGN1-EGFP. GCs that coexpressed NLGN1 and ASTN2 had higher levels of internalized NLGN1 after a 20 min chase than GCs that expressed NLGN1 and a control plasmid (Fig. 3c, d). Thus, ASTN2 interacts with several key synaptic adhesion proteins and can reduce their surface expression in neurons and HEK cells by endocytosis.

### ASTN2 binds a number of proteins suggestive of trafficking of multiple protein complexes in neurons

To identify additional ASTN2 binding partners in an unbiased manner, we carried out IP of ASTN2 from the juvenile (P22-28) mouse cerebellum followed by mass spec analysis. An initial round of experiments with duplicate samples was followed by a second experiment with more stringent washes and the inclusion of a further negative control in which the ASTN2 antibody was affinity removed from the antisera (Fig. S6a). The combined experiments identified 466 proteins enriched in the ASTN2 IP compared to IgG or the depleted ASTN2 sera samples (Table S2). Further refinement of the list to only include proteins with at least 3 peptide hits that were ?1.5 fold enriched in the ASTN2 IP versus the IgG or the depleted anti-ASTN2 sera yielded 57 proteins (Fig. 4a). We identified AP-2, an adaptor protein in Clathrin-mediated endocytosis from the plasma membrane, and VPS36, found on sorting endosomes. We also identified multiple proteins involved in synaptic form and function such as C1q, shown to mediate synaptic pruning (18), OLFM1/3, which form complexes with AMPA receptors (19) and were recently identified as ASD candidates (20), ROCK2, a Rho-kinase which regulates spine morphology and synaptic activity through regulation of the cytoskeleton (21), and SLC12a5 (KCC2), a potassium/chloride co-transporter that regulates the intracellular Chloride ion gradient as well as dendritic spine morphogenesis (22, 23), and is also implicated in ASD (24–26). Thus, ASTN2 interacts with multiple proteins that regulate synaptic activity. We confirmed these interactions by co-IP and Western blot in HEK 293T cells (Fig. 4c, Fig. S6b). Furthermore we detected co-IP of ASTN2/ROCK2/AP2 (Fig. 4d) and ASTN2/NGLN2/AP2 (Fig. S6c) *in vivo.* Functional enrichment analysis of the proteins identified categories (Fig. 4b) such as phagosome, endocytosis, synapse, microtubule-associated processes, and t-RNA splicing ligase complex (not investigated further here). A number of the interacting proteins identified are, like ASTN2, implicated in ASD pathogenesis (asterisks, Fig. 4b). Taken together, our IP/mass spec experiments show that ASTN2 binds to proteins involved in vesicle trafficking and synaptic function including synaptic pruning proteins, ion transporters, accessory proteins to ligand-gated ion channels, and proteins involved in cytoskeletal rearrangements, suggesting that ASTN2 possibly promotes the trafficking of multiple surface proteins.

**Figure 4.**
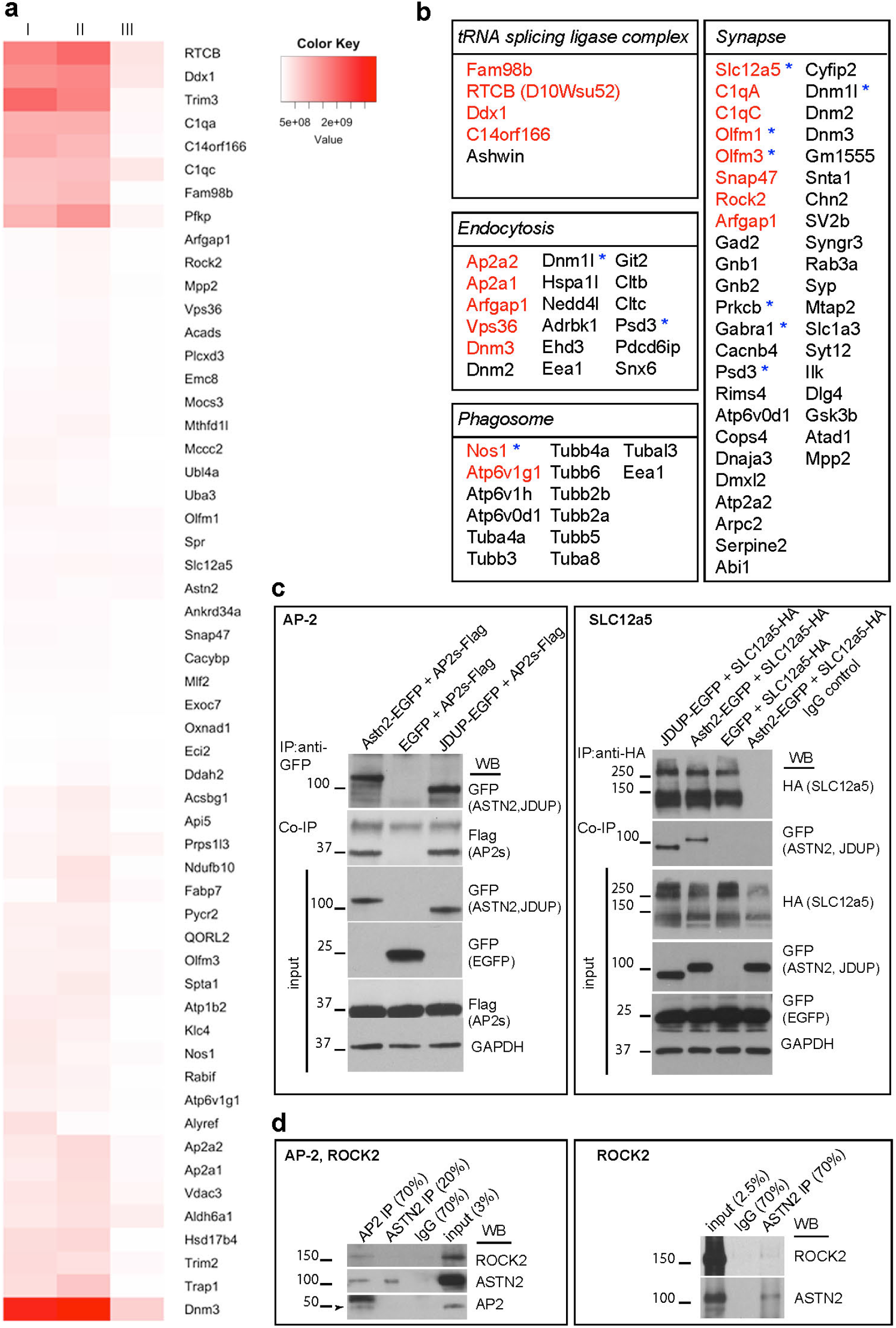
Protein interactors of ASTN2 identified by IP plus LC-MS/MS. (**a**) Heat map of the top 57 candidate interacting proteins enriched in ASTN2 IPs in three experiments from P22-28 cerebellar lysates. The intensity of the map is based on the MS intensity spectra values (see Table S2). IP I and II are biological and technical replicates. IP III is a third biological replicate, which was washed more stringently and performed separately (see methods). **(b)** Lists of identified proteins by functional enrichment. Proteins belonging to the list of top hits (in a) are shown in red. ASD-associated proteins found by cross-referencing Table S2 to the SFARI human ASD-gene list (www.gene.sfari.org) are marked with blue asterisks. **(c, d)** Western blots showing co-IPs of AP-2 (sigma fragment, Ap2s) and SLC12a5 with ASTN2 or JDUP in HEK 293T cells (c) and of AP2, ROCK2, and ASTN2 in cerebellar lysates at P22 (d). Protein ladder in kDa. In the SLC12a5 blot GFP appears in all samples due to the existence of an IRES-EGFP in the SLC12a5-HA construct.

### ASTN2, but not the FNIII truncation, induces degradation of surface proteins

In our flow cytometry analyses (Fig. 3b), an increase in the percentage of cells that did not express NLGN1 or 3 in the presence of ASTN2 was observed (bottom left quadrants of graphs), as opposed to an increase in cells that expressed NLGN1/3 internally but not on the cell surface (bottom right quadrants), suggesting that ASTN2 not only internalizes proteins but also induces degradation. Indeed, we detected reduced expression of the identified synaptic binding partners (NLGN1-4, SLC12a5, OLFM1, Fig. 5a, b), but no change in the levels of the adaptor protein AP-2 or GAPDH (Fig. 4c, 5a, b) upon ASTN2 overexpression compared to controls by Western blot. Importantly, while co-expression of NLGN1 or SLC12a5 with ASTN2 resulted in reduced levels of both, this reduction was much less marked upon coexpression with JDUP (Fig. 5b). Thus, co-expression of ASTN2 but not the JDUP truncation markedly reduced protein levels. Moreover, shRNA-mediated knockdown of ASTN2 resulted in similar levels of NLGN1 and SLC12a5 to cells without ASTN2 protein or cells with JDUP, suggesting that the reduction the levels of ASTN2 binding partners only occurs in the presence of intact ASTN2. Together, our data suggest that ASTN2 promotes the internalization and degradation of surface proteins.

**Figure 5.**
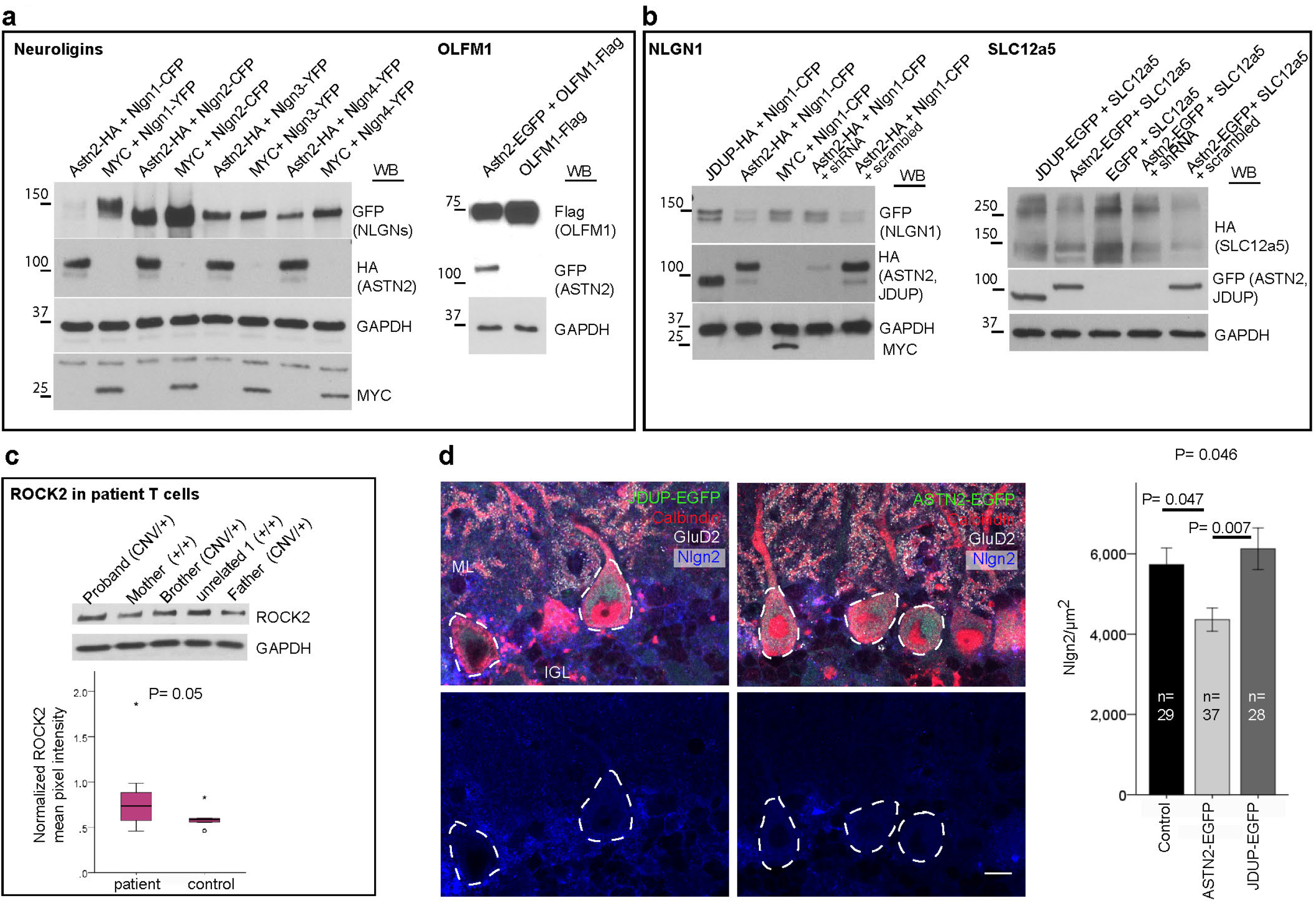
ASTN2 reduces the levels of interacting proteins. (**a**) Western blots showing reduced protein levels of NLGN1-4 and OLFM1 in HEK 293T cells in the presence of ASTN2 as compared to MYC (control) or OLFM1 alone. (**b**) Western blots showing reduced expression of NLGN1 and SLC12a5 in HEK 293T cells in the presence of ASTN2 or ASTN2 co-expressed with a scrambled plasmid, but less so in the presence of JDUP, MYC, or when ASTN2 is knocked-down with shRNA. GAPDH was used as an internal control for protein loading. **(c)** A representative Western blot showing ROCK2 levels in *ASTN2* CNV patient T cells. The controls consisted of the mother and unrelated healthy subjects. Quantification of ASTN2 normalized to GAPDH in four technical replicates of 3 patients and 3 controls is shown in the box plot. **(d)** Conditional expression of ASTN2-EGFP and JDUP-EGFP (green) in sagittal sections of *PCP2-Cre+* cerebella labeled with antibodies against Calbindin (red), NLGN2 (blue) and GluD2 (white). Quantification of NLGN2 levels (corrected integrated pixel density) in PC somas (outlined by dashed lines) upon ASTN2 versus JDUP overexpression or control (*PCP2-Cre+* mice injected with conditional ASTN2-EGFP virus). Graph shows means +/−1 SEM. n= total number of cells analysed from 3 mice per condition. P-value at top by ANCOVA and closer to bars by post-hoc tests between groups. Scale bar: 10μm

To further examine the idea that ASTN2 promotes protein degradation, we searched the top 57 protein hits identified by mass spec to see if any membrane proteins identified were also found in CD4+ patient T-cells, where ASTN2 levels are reduced. Among the 57, only ROCK2 is expressed in neurons as well as in T-cells. Although like ASTN2, ROCK2 levels were variable among patients, they were on average higher in patients compared to controls (Fig. 5c). The expression level of the lower ASTN2 band (Fig. 1c bottom right graph) inversely correlated with ROCK2 levels among patients. Together, our data suggest that ASTN2 plays a fundamental role in modulating the dynamic localization and degradation of several protein complexes in multiple cell types.

### ASTN2 modulates synaptic activity

To investigate whether manipulation of ASTN2 levels impacts synaptic function in the cerebellum, we generated conditional lentiviruses expressing either the full length EGFP-tagged ASTN2 (pFU-cASTN2-EGFP) or a truncated version (pFU-cJDUP-EGFP) lacking the FNIII domain. Overexpression approaches have generally been reported to be more sensitive in revealing roles for adhesion molecules (27) and proteins in multimeric complexes during synaptogenesis, due to the disruption of the stoichiometry of complexes and unmasking of functions otherwise compensated for by homologous proteins in loss of function approaches (28). Viruses were stereotactically injected *in vivo* into *Pcp2-Cre+* cerebella at P0-2 (Fig. 6a) to target PCs. EGFP expression, restricted by *Pcp2-Cre* to PCs only, was observed 3-4 weeks after viral injection (Fig. 6b). Interestingly, ASTN2-EGFP, but not JDUP-EGFP expression, resulted in mislocalization of some PCs to ectopic locations within the internal granule cell layer and in the white matter (Fig. 6b and Fig. S7).

**Figure 6.**
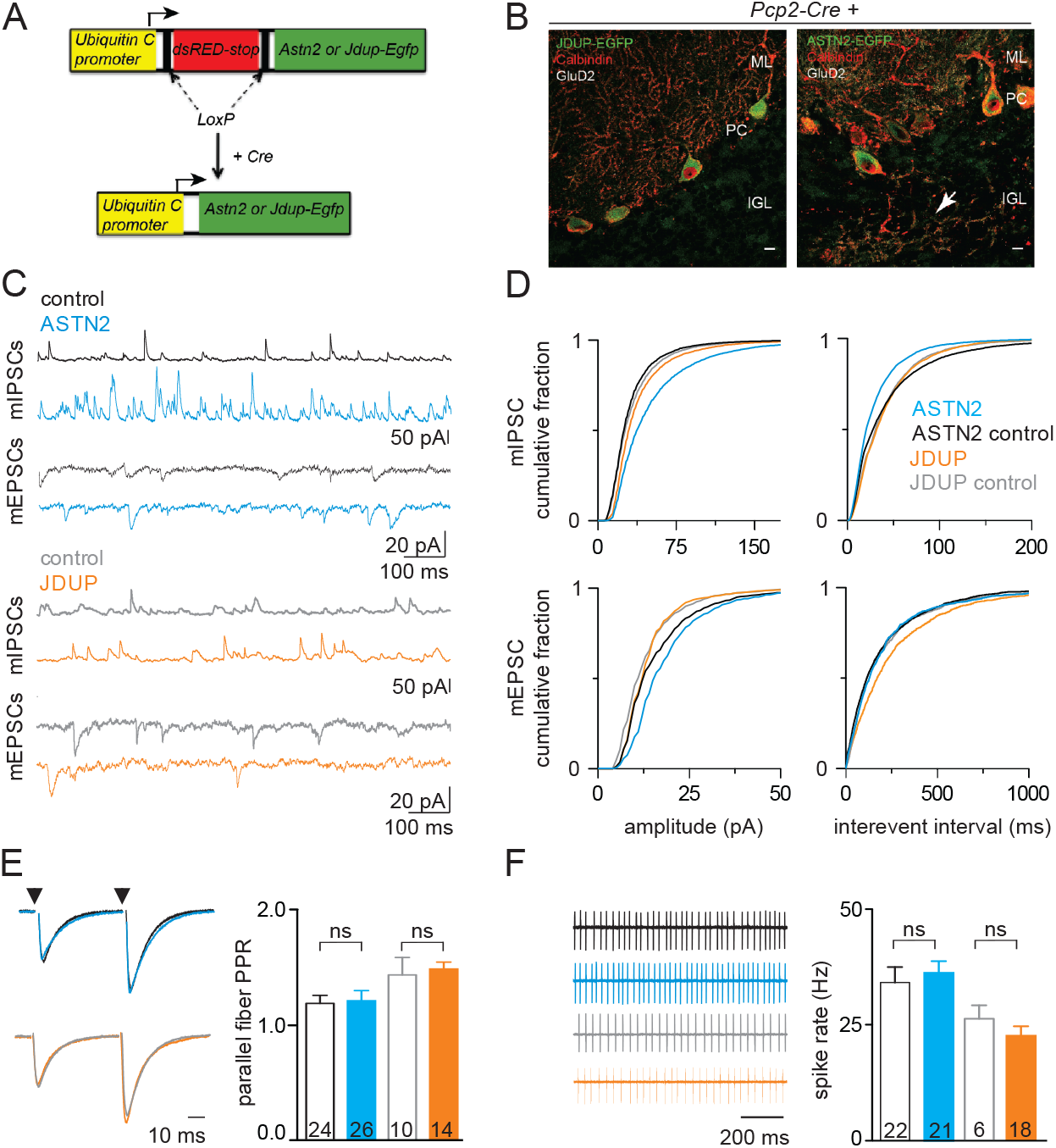
Effect of ASTN2 overexpression on synaptic activity of Purkinje cells. **(a)** Schematic of conditional lentiviral vectors. Expression of ASTN2-EGFP or JDUP-EGFP is driven by the Ubiquitin C promoter in the presence of Cre. **(b)** Sagittal sections showing JDUP-EGFP and ASTN2-EGFP (green) expression in PCs marked by Calbindin (red) and GluD2 (white) 3-4 weeks after injection into *Pcp2-Cre+* mice. Arrow indicates ectopic PC in the IGL of an ASTN2-EGFP injected mouse. **(c)** Miniature inhibitory (mIPSCs, top) and excitatory (mEPSCs, bottom) postsynaptic currents in control (*PCP2-Cre-^/−^*; black, n=21 cells) and ASTN2 expressing PCs (*PCP2-Cre+;* blue, n=14 cells) and in control (*PCP2-Cre-^/−^*; gray, n= 10 cells) and JDUP expressing PCs (*PCP2-Cre+;* orange, n= 12 cells). **(d)** Cumulative histograms of the amplitude (left) and frequency (right) miniature events in control (black or gray), and ASTN2 (blue) or JDUP (orange) expressing PCs. Distributions were compared using the Mann-Whitney U test and were found significantly different between ASTN2 and controls in all measurements (P<0.0001) except for mEPSC frequency which was the same between control and ASTN2, but significantly different between control and JDUP (P<0.0001). **(e)** Left: Evoked parallel fiber EPSCs (Vm ∼ −75 mV; 50 ms inter-stimulus interval, arrowheads). Right: summary graphs of paired-pulse ratios (mean +/−1 SEM) **(f)** Left: cell-attached recordings of spontaneous spiking. Right: summary graphs (mean +/−1 SEM) of spontaneous firing rates. n= total number of cells recorded from 5-7 animals per condition. Scale bars: 10μm. IGL, internal granule layer; ML, molecular layer; PC, Purkinje cell layer; ns= not significant.

To test the properties of intrinsic excitability and synaptic transmission onto PCs that expressed either ASTN2-EGFP or JDUP-EGFP, we performed whole-cell electrophysiological recordings in acute brain slices from injected *Pcp2-Cre+* animals 3-4 weeks after viral injections (P21-35). Control recordings were performed on EGFP-negative PCs from *Pcp2-Cre^−/−^* littermates injected with the same conditional viruses. Miniature excitatory/inhibitory postsynaptic currents (mEPSCs/mIPSCs) were recorded to assess non-evoked, quantal synaptic input, primarily from the parallel fibers (mEPSCs) and the inhibitory stellate and basket cells (mIPSCs). In PCs expressing ASTN2-EGFP, we found a significant increase in both mIPSC amplitude (Δ_max_ =25.2%) and frequency (Δ_max_=13.8%) and an increase in mEPSC amplitude (Δ_max_ =21.5%) in the same PCs. There was no change in the frequency of mEPSCs (Δ_max_ =5.9%, Fig. 6c, d). In PCs expressing JDUP-EGFP, there was a much less marked increase in mIPSC amplitudes (Fig. 6c, d, Δ_max_=11.4%) and no change in frequency (Δ_max_ =2.3%). In addition, we observed a less marked increase in mEPSC amplitudes (Δ_max_=11.9%) but a significant decrease in mEPSC frequency (Δ_max_=13.5%). These results indicate changes in the synaptic strength of PCs, with the strongest effect on mIPSCs upon ASTN2-EGFP expression.

We also tested evoked excitation from parallel fibers, and found that the paired pulse ratio was unchanged, suggesting no difference in presynaptic release probability (Fig. 6e). In addition, we did not observe any differences in the spontaneous spiking of either ASTN2-EGFP or JDUP-EGFP expressing cells as measured by non-invasive cell-attached recordings (Fig. 6f). These results suggest that ASTN2 overexpression increases synaptic strength primarily by altering the properties of the postsynaptic membrane, rather than the intrinsic excitability of PCs or presynaptic release dynamics. Importantly, we did not observe the same degree of changes with expression of JDUP-EGFP as we did with ASTN2-EGFP. Finally, comparison of NLGN2 expression, which is the most highly expressed of the Neuroligins in the cerebellum (29), in the soma of targeted PCs revealed a significant decrease in ASTN2-EGFP PCs compared to JDUP-EGFP or control PCs from *Pcp2-Cre^−/−^* cerebella injected with the same conditional viruses (Fig. 5d), further corroborating our earlier findings that ASTN2 overexpression induces degradation of synaptic binding partners.

## Discussion

Using immuno-gold EM, biochemical, electrophysiological, and functional assays we demonstrate a novel role for ASTN2 in controlling protein trafficking and homeostasis in synaptic function. We detect ASTN2 in dendritic spines of neurons and in trafficking vesicles, and identify binding to several synaptic as well as trafficking proteins. Consistent with this interpretation, ASTN2 binds the Clathrin adapter AP-2. In postmitotic PCs, the sole output neuron of the cerebellar cortex and a cell type with documented loss in ASD patients (14), overexpression of ASTN2 increased synaptic strength and decreased protein levels of synaptic binding partners. Our analyses suggest a role for ASTN2 in controlling surface membrane protein dynamics and underscore the contribution of impairment in protein trafficking to neurodevelopmental brain disorders.

The results reported here compared the effect of ASTN2 overexpression with that of a truncated form lacking the FNIII domain. These experiments were informative in showing that removal of the FNIII domain interfered with the ability of ASTN2 to promote protein degradation, but not its ability to interact with binding partners. Overexpression in cell types that normally express ASTN2 provided a powerful means to study the consequences of disrupting the stoichiometry of ASTN2 in its native protein complexes. It should be noted that knock-down of ASTN2 protein was achieved in HEK cells but not in neurons (Fig. S3) possibly due to the formation of protein complexes in neurons, which promote the perdurance of the protein. Indeed, two pulse-chase studies examining protein turnover in the mouse brain found that ASTN1, the homologue of ASTN2, is present even after 1 month following *in vivo* isotopic labeling (17, 30). It is therefore likely that ASTN2 is also extremely long-lived.

The interpretation that ASTN2 is a long-lived protein is also consistent with the reported stability of ASTN2 protein at pH 4.0 (31), which would allow it to traffic binding partners through the lower pH endosomal compartments of the endo/lysosomal system. In line with a trafficking role, the ASTN2 protein sequence (Fig. S2) contains tyrosine-based sorting signals recognized by the adaptor proteins AP1-4, and a dileucine-based signal recognized by Golgi-localized Y-ear containing ARF-binding (GGA) proteins (localized to endosomes), also involved in endosomal trafficking (32), as well as lysosomal sorting signals like those found in lysosomal membrane proteins LAMP1/LAMP2. These signals are not only recognized at the plasma membrane, but also at sorting stations such as the endosomes (32), suggesting that ASTN2 is likely involved in multiple steps of endo/lysosomal trafficking and not just at the surface membrane. This is consistent with our EM data, which showed that ASTN2 localizes to membranes and vesicles throughout the cell soma (Fig. 2d), and with our co-localization experiment with endosomal markers where ASTN2 was found in a small fraction of early as well as late endosomes (Fig. S4). Furthermore, our mass spec and biochemical data show that ASTN2 binds various endosomal trafficking and sorting proteins including AP-2 and VPS36 (part of the ESCRTII complex) and controls the surface removal and degradation of a number of synaptic proteins. Finally, our EM analysis showed ASTN2 also on autophagosomes. As a large body of work shows that autophagic vesicles fuse with endosomes and lysosomes, (33, 34), it will be interesting to further examine the involvement of ASTN2 in the interplay between autophagy and endo/lysosomal trafficking. We note that our findings do not exclude a role for ASTN2 in protein degradation pathways other than the endo/lysosomal system.

In addition to components of vesicle trafficking, the identification of interactions with multiple proteins involved in synaptic pruning, C1q (18), AMPA receptor accessory proteins, OLFM1/3 (19), ion transport, SLC12a5 (22), and proteins regulating synaptic adhesion and activity, ROCK2 (21) and NLGN1-4 (29), suggest that ASTN2 may modulate the composition of multiple protein complexes which impact synaptic form and function. In PCs, the sum of these interactions is increased postsynaptic activity upon ASTN2 overexpression. We speculate that the ability of ASTN2 to remove surface proteins also caused the mislocalization of PCs *in vivo* (Fig. S7). The mechanisms that keep PCs in place however are not fully understood.

Of note, a previous proteomic study of synaptosomal fractions prepared from mouse and human brains detected ASTN2 (35), corroborating our finding that ASTN2 is indeed present near synapses. The largest synaptic activity changes observed upon ASTN2 overexpression were increased mIPSC frequency and amplitude. Increases in mini frequency are commonly associated with increased numbers of synapses, while increases in mini amplitude often reflect increased numbers of postsynaptic receptors. However, other mechanisms could also contribute, such as alterations in single channel conductance, receptor desensitization, changes in intracellular ion concentrations due to alterations in plasma membrane ion transporters, or fine-scale structural changes (36). Given the finding that ASTN2 internalizes synaptic and cell surface proteins, we favor the second set of scenarios whereby removal of proteins accessory to channels, such as OLFM1/3 and NLGN2, or the ion transporter SLC12a5, change ligand-gated channel mediated responses. Excessive removal and degradation of accessory proteins could also leave receptors stranded on the cell surface, with their normal activity disrupted. Moreover, reduced levels of SLC12a5 could result in an increase in both mEPSCs and mIPSCs as reported by a study in the hippocampus of SLC12a5 deficient mice (37). It should be noted that although NLGNs were not identified by IP/mass spec, which samples the most abundant and stable interactions, two NLGN1/3 peptides were identified with a targeted mass spec approach (data not shown). It is possible that other transient interactions were not detected by our method and would need to be investigated in a candidate protein approach. Overall, our data suggest that as the primary role of ASTN2 appears to be in trafficking of an array of synaptic proteins, manipulations of ASTN2 can result in diverse synaptic modifications including changes in postsynaptic receptor expression and synaptic strength depending on context. Further studies on the effect of loss of ASTN2 await development of a genetic mouse model.

Interestingly, the JDUP truncation did not abolish, but rather altered the interaction of ASTN2 with its binding partners, increasing its affinity for some (NLGN3/4) while decreasing it for others (NLGN1/2, Fig. 3a). These differences in affinities could underlie the differing effects on mIPSC and mEPSC events induced by ASTN2 versus JDUP (Fig. 6). While ASTN2 clearly interacts with and regulates the availability of a number of proteins, a systems level analysis using advanced live cell imaging combined with single molecule tracking of multiple proteins in action is needed to understand the combined effects of manipulation of an ASTN2-mediated trafficking pathway.

As reported here and elsewhere, patients with *ASTN2* CNVs can manifest a spectrum of NDD phenotypes, even within the same family (Table S1). Our analysis of the association of *ASTN2* CNVs with specific features commonly reported in such patients, revealed a high level of association with ID and delayed speech and language. Based on our findings that an ASTN2-mediated protein trafficking pathway modulates synaptic strength, we propose that *ASTN2* CNVs (truncating duplications and deletions) in NDD patients, cause an accumulation of surface proteins. It appears critical for neurons to respond to inputs and modify their surface proteome in a rapid fashion. Activity-dependent changes in gene expression have been well documented, but more rapid changes in the composition of the synaptic proteome, through protein trafficking and degradation, would indeed offer quicker means to adjust to such inputs. Interestingly, *Astn2* levels have been reported to change in response to stress in the CA3 region of the hippocampus in mice (38), suggesting the possibility that ASTN2 may modulate the surface proteome in response to activity in neurons.

ASTN2 is unusual in that it is so abundantly expressed in the cerebellum compared with other brain regions. Although the contribution of the cerebellum to ASD, ID, or speech and language development is poorly understood, neuroimaging studies show both functional and neuroanatomical evidence for the critical importance of the cerebellum (39–41). Computational studies mapping the spatio-temporal co-expression of ASD-associated genes, including ASTN2, show that cortical projection neurons (layer 5/6) and the cerebellar cortex are the two most prevalent sites of ASD gene co-expression (42, 43). Furthermore, long term depression (LTD), which is thought to be essential for many forms of cerebellar learning, is altered at parallel fiber-PC synapses in mice with targeted disruptions of several ASD-associated genes (16, 44–47). The present findings suggest that ASTN2 is a key regulator of dynamic trafficking of synaptic proteins in the cerebellum and lend support to the idea that aberrant regulation of protein homeostasis is a contributing cause of complex neurodevelopmental disorders such as ASD and ID (48).

## Methods

### Human subjects

Patient clinical information and PBMCs were collected at Kennedy Krieger Institute (Baltimore) upon informed consent from all subjects. This study was approved by IRB boards of the Kennedy Krieger Institute (Baltimore) and The Rockefeller University (New York).

### Mice

C57Bl/6J mice (Jackson Laboratory) were used unless stated otherwise. All procedures were performed according to guidelines approved by the Rockefeller University Institutional Animal Care and Use Committee. Both males and females were used for all studies and were randomly allocated to control and test groups. *Reverse transcription polymerase chain reaction (RT-PCR) and quantitative (q) RT-PCR:* mRNA was extracted using the AllPrep DNA/RNA/Protein Mini Kit (Qiagen) and cDNA transcribed with the Transcription First Strand cDNA Synthesis Kit (Roche) according to the manufacturer’s description. RT-PCR and qRT-PCR were carried out according to standard procedures. See supplementary methods.

### Primary cell and cell line culture

HEK 293T/17 cells (ATCC#Crl-11268) were grown at 37°C/5% CO_2_ in DMEM/F12, 10% heat inactivated Fetal Bovine Serum (FBS), 4mM L-glutamine, 100U/ml Penicillin/Streptomycin (all from Gibco). Mixed cerebellar cultures were prepared from P6-8 pups and cultured in serum containing medium as previously described (1, 49). Half of the medium was replaced with fresh medium without serum every 3-4 days for the duration of culture. Both HEK cells and primary granule cells (at DIV14) were transfected using Lipofectamine 2000 (Invitrogen) according to the manufacturer’s description. PBMCs from human subjects were isolated using BD Vacutainer™ CPT™ Tubes (BD Biosciences #362753). 10 × 10^6 cells were first plated in T cell medium (RPMI-1640, 10% FBS, 2 mM Glutamine, 100 U/ml Penicillin/Streptomycin, 10 μM HEPES) for 3 hrs, allowing monocytes to attach to the plate and be discarded. CD4+ cells were then sorted by MACS Separation using CD4 MicroBeads (human) and MS columns (Miltenyi Biotec) according to the manufacturer’s descriptions and expanded in T cell medium containing 30 U/ml IL2 (Peprotech #200-02) and CD3/CD28 beads (Dynabeads #11161D) according to the manufacturer’s description. Media change (T cell medium containing IL2) was performed every two days and cells were collected at DIV 8 for mRNA and protein analysis.

### Immunohisto/cytochemistry

Mice P15 or older were fixed by perfusion with 4% PFA and sectioned sagittally at 50μm (Leica Vibratome). *In vitro* cultured cells (described above) were grown on glass coverslips (no 1.5 thickness, Fisher Scientific) and fixed for 15 minutes at room temperature in 4% PFA. Immunohistochemistry was carried out according to standard protocols. See supplementary methods.

### Antibodies used for immunohistochemistry, immunoprecipitation and Western blot

see supplementary methods.

### cDNA/ShRNA constructs

see supplementary methods.

### Knockdown of ASTN2

HEK cells were transfected, as described earlier, with plasmids expressing the *Astn2* cDNA alone or together with shRNA and scrambled constructs. Two days later, cells were processed for immunohistochemistry and Western blot as described earlier. For knockdown in neurons see supplementary methods.

### Pre-embedding nanogold immunolabeling and electron microscopy

P28 mice were perfusion fixed with 4% PFA. 50μm sagittal vibratome sections were prepared and incubated in blocking solution (3% BSA, 0.1% saponin in 0.1M sodium cacodylate buffer, pH7.4) for 2 hr at room temperature, followed by incubation in anti-ASTN2 antibody in blocking solution for 48 hrs at 4°C. After washing (4x 1 hr in sodium cacodylate buffer), the sections were incubated for 2 hrs at room temperature with secondary antibody (1:100, Nanoprobes-Nanogold, anti-Rabbit: 2003), washed 4x 1 hr in 0.1% saponin in 0.1M sodium cacodylate buffer, pH 7.4 and then fixed in 2.5% glutaraldehyde (Sigma) overnight at 4°C. The sections underwent silver enhancement (HQ Silver Enhancement 2012, Nanoprobes), and Gold Toning using a 0. 1% solution of gold chloride (HT1004, Sigma) according to the manufacturer’s description. The tissue was post-fixed with 1% osmium tetroxide for 1 hour on ice. Sections underwent en bloc staining with 1% uranyl acetate for 30 min, dehydrated in a graded series of ethanol, incubated for 10 minutes in acetone, infiltrated with Eponate 12™ Embedding Kit (Ted Pella), and polymerized for 48 hrs at 60°C. 70nm ultrathin sections were imaged on a JEOL JEM-100CX at 80kV and a digital imaging system (XR41-C, Advanced Microscopy Technology Corp, Woburn, MA). For negative controls, sections were processed as described but with omission of the primary antibody.

### Immunoprecipitation, intracellular crosslinking of proteins, depletion of ASTN2 antisera and Western blot

For *in vitro* IPs, proteins from transfected HEK 293T cells were extracted in either RIPA buffer (Thermo Scientific) or in a customized IP buffer (50 mM Tris pH 7.4, 200 mM NaCl, 5 mM MgCl2, 5 mM NaF, 1.5% Octyl β-D-glucopyranoside (abcam), 1x Protease Inhibitor Cocktail, 10U Benzonase Nuclease, Sigma). For *in vivo* IPs, proteins were extracted from P22-28 cerebella using the customized IP buffer (above). 1.8 mg (*in vivo*) or 0.5 mg (*in vitro*) protein inputs were used to carry out overnight IPs according to standard protocols using Dynabeads Protein G (Invitrogen) cross-linked with antibodies using Bis-sulfosuccinimidyl suberate (BS3) cross-linking according to the manufacturer’s description (Thermo Scientific). As controls, either normal IgG from the same species as the antibody (Santa Cruz) or an “anti-ASTN2 depleted antisera” was used. The anti-ASTN2 depleted antisera was prepared by incubating the anti-ASTN2 antibody with N-terminal biotinylated peptide against which the antibody had been raised (KITCEEKMVSMARNTYGETKGR) in the customized IP buffer. The antibody-peptide mix was then pulled out with Dynabeads MyOne Streptavidin T1 beads (Invitrogen) and the volume containing the antisera depleted of ASTN2 antibody was cross-linked to Dynabeads Protein G for use as negative control. Depletion of anti-ASTN2 antibody was confirmed by Western blot analysis on cerebellar lysates (Figure S6a). Intracellular cross-linking of proteins was carried out prior to some IPs (as indicated in main text) with disuccinimidyl suberate DSS (Thermo Scientific) according to the manufacturer’s descriptions. Western blots were carried out according to standard protocols using SDS-PAGE gels (Fisher) and Immobilon-P transfer membranes (Millipore). Blots were developed using an ECL Western Blotting Kit (GE Healthcare) or SuperSignal West Pico kit (Thermo Scientific) and exposed to X-ray film (Kodak).

### Mass spectrometry

IPs were prepared as described earlier using 1.8 mg protein input from whole cerebellar lysates (P22-28) with antibody cross-linked beads. Immuno-precipitated proteins were eluted with 8M urea (GE Healthcare) in 0.1 M ammonium bicarbonate (Sigma) and 10 mM DTT (Sigma). Cysteines were alkylated with iodoacetamide (Sigma). Samples were then diluted below 4 M urea before digesting with LysC (Waco Chemicals) for 6 hrs, after which urea was diluted below 2 M for overnight trypsin (Promega) digestion. Peptides were desalted by StaGE tips and processed for nano LC-MS/MS in data-dependent mode (Dionex U3000 coupled to a QExactive or QExactive Plus mass spectrometer, ThermoFisher Scientific). Generated LC-MS/MS data were queried against Uniprot’s complete Mouse Proteome (downloaded July 2014) concatenated with common contaminants, and peptides were identified and quantified using Proteome Discoverer 1.4 (ThermoScientific) and Mascot 2.5.1 (Matrix Sciences) with fully tryptic restraints (Trypsin/P) and up to 3 missed cleavages with Protein N-term acetylation and methionine oxidation as variable modifications and cysteine carbamidomethylation as a stable modification. Peptide matches required 5 ppm accuracy in MS1 and 20 mmu in MS2, with a 1% FDR filter using Percolator (55). To create the list of 466 identified proteins, all proteins that were only present in IgG samples were filtered out as were proteins that showed enrichment in the IgG control over the ASTN2 IP in the most stringently washed experiment (III). The most stringent list of 57 proteins (Fig. 4) was created by including only proteins with at least 3 peptide hits that were ?1.5 fold enriched in the combined ASTN2 I+II IPs/ I+II IgG IPs or in ASTN2 III IP/ depleted anti-ASTN2 sera. Functional enrichment analysis was carried out using STRING (v10.0, http://string-db.org/).

### Flow cytometry

Transfected HEK 293T cells were harvested in 1 mM EDTA in PBS. The surface fraction of GFP-linked surface proteins was immuno-labeled (live) with rabbit anti-GFP followed by Alexa-647 anti-rabbit and cells were stained with Propidium Iodide (Sigma-Aldrich) for dead cell exclusion. Flow cytometry analysis (BD Accuri C6, BD Biosciences) was carried out using 488 nm and 640 nm lasers and the CFlow Sampler software (BD Biosciences). A total of 20,000 single viable cells, identified by size and lack of Propidium Iodide staining, were analyzed per condition (resulted in approximately 100,000 events per condition). Gates were set using non-transfected control cells and cells expressing cytosolic GFP (α-tubulin-GFP), which were processed for live GFP labeling as described above. Data were analyzed by FlowJo v.9.3.3 (TreeStar Inc., Ashland, OR).

### Surface and pulse-chase labeling in neurons

DIV14 mixed cerebellar cultures (prepared as described earlier) were transfected with either “Nlgn1-HA-YFP + Astn2-EGFP” or “Nlgn1-HA-YFP + EGFP” plasmids using Liopefectamine 2000. On DIV17, cells were incubated with anti-GFP (1:500) diluted in culture medium containing 10 mM HEPES (Sigma) at 4°C for 20 min, followed by 2x washes with medium + 10 mM HEPES on ice to prevent endocytosis. For surface labeling experiments (Fig. S5), cells were fixed and processed as previously described. For pulse-chase experiments (Fig. 3) cells were incubated in fresh medium for a 20 min chase period at 35°C/5% CO_2_. Cells were then incubated with anti-rabbit Alexa-633 (1:300) for 30 min at 4°C to label all pulsed NLGN1-HA-YFP left on the cell surface, followed by 2x washes in medium on ice to wash away unbound antibodies. The cells were then fixed and processed as described earlier to detect internalized (antirabbit Alexa-555 secondary) and total (mouse anti-HA primary followed by antimouse Alexa 405 secondary) protein. Control experiments were carried out to ensure that surface labeling did not occur with the GFP antibody on cells that expressed EGFP or ASTN2-EGFP only, as neither protein is exposed on the surface membrane, while the YFP tag of NLGN1-HA-YFP is positioned outside the plasma membrane and hence detected by live labeling.

### Imaging

Images were acquired using an inverted Zeiss LSM 880 NLO laser scanning confocal microscope with a Plan-Apochromat 40x/1.4 NA objective oil immersion lens and 2.8 x digital zoom. For the lower power image in Fig. 2a a Plan-Apochromat 10x/0.45 NA lens was used. Images were acquired by setting the same gain and offset thresholds for all images per experiment and over/underexposure of signal was avoided. Images were quantified in ImageJ (version 2.0.0-rc-38/1.50b) unless stated otherwise.

### Virus production and in vivo viral injections

VSV-G pseudo-typed lentiviruses were produced with the pFU-cASTN2-EGFP and pfU-cJDUP-EGFP plasmids as previously reported (54). Viruses were collected and concentrated 45 hrs post transfection and the pH of the media was kept between 7-7.3. Neonatal (24-30 hrs old) F1s from hemizygous PCP2-Cre breeding pairs (Jackson Laboratory, B6.Cg-Tg(Pcp2-cre)3555Jdhu/J) were cryoanesthetized and injected using a modified protocol of Kim et al. (56) using a 10 μL Hamilton syringe (Hamilton #1701-RN) fitted with a custom 32 gauge needle (point style: #4; angle: 12°; length: 9.52 mm; Hamilton #7803-04). The needle was inserted perpendicular to the occipital plate at a depth of ∼2.5 mm, centering the tip in-line with the anterior-posterior axis and between the ears. All procedures were approved by the Duke University Institutional Animal Care and Use Committee and were in compliance with regulations.

### Electrophysiology

Acute sagittal slices (250 μm thick) were prepared from the cerebellar vermis of 3-4 week old injected (PCP2-Cre+) and control littermates (PCP2-Cre-/−). Slices were cut in an ice-cold potassium cutting solution (57) consisting of (in mM): 130 K-gluconate, 15 KCl, 0.05 EGTA, 20 HEPES, 25 glucose, pH 7.4 with KOH, and were transferred to an incubation chamber containing artificial CSF comprised of (in mM): 125 NaCl, 26 NaHCO3, 1.25 NaH2PO4, 2.5 KCl, 2 CaCl2, 1 MgCl2, and 25 glucose (pH 7.3, osmolarity 310). Electrophysiological recordings were performed at 32-33° C using a Multiclamp 700B amplifier (Axon Instruments), with signals digitized at 50 kHz and filtered at 10 kHz. All whole-cell recordings were performed using a cesium-based internal solution containing (in mM): 140 Cs-gluconate, 15 HEPES, 0.5 EGTA, 2 TEA-Cl, 2 MgATP, 0.3 NaGTP, 10 Phosphocreatine-Tris2, 2 QX 314-Cl. pH was adjusted to 7.2 with CsOH. For parallel fiber stimulation experiments, glass monopolar electrodes (2-3 MΩ) were filled with aCSF, and current was generated using a stimulus isolation unit (A.M.P.I., ISO-Flex). Spontaneous miniature synaptic currents were recorded in the presence of tetrodotoxin (TTX, 0.5 μM, Tocris). IPSCs were recorded at the empirically determined EPSC reversal potential (∼+10 mV), and EPSCs were recorded at the IPSC reversal potential (∼+75 mV). Membrane potentials were not corrected for the liquid junction potential. Series resistance was monitored with a −5 mV hyperpolarizing pulse, and only recordings that remained stable over the period of data collection were used. Miniature IPSCs and EPSCs were analyzed using MiniAnalysis software (v6.0.3, Synaptosoft Inc.), using a 1 kHz low pass Butterworth filter and a detection threshold set to 5x (for IPSCs) or 10x (for EPSCs) higher than baseline noise. So that no individual recording biased our distributions, 400 mIPSCs and 120 mEPSCs from each cell were randomly selected to establish the amplitude and frequency distributions of events across conditions. To measure the paired pulse ratio, parallel fibers were stimulated at 20 Hz.

### Quantification and statistics

Observations were replicated in at least three independent experiments (technical replicates). Data represented in graphs are both biological (pooled or individual animals/starting material) and technical (repeated multiple times) replicates, except for Fig. 1c, where individual data points are shown for human samples. Pixel intensities of Western Blots and immunolabelings were quantified using ImageJ. Surface and internal labeling of NLG1-HA-YFP as well as NLGN2 labeling in PCs *in vivo* were quantified as follows: each cell and its processes including dendritic spines in the case of GCs, and the cell soma only in the case of PCs *in vivo*, were outlined. The “integrated density” was measured (sum of all pixel intensities/μm2). The “mean fluorescence background” of each channel was also measured by selecting an area containing no cells. The “corrected fluorescence” was then calculated per cell as: integrated density – (area of selected cell * mean fluorescence intensity of image). For GCs, 20 cells per coverslip and 2 coverslips per condition were imaged from three independent experiments. The data plotted as mean+/−1 SEM. “total” in Fig. 5d represents the sum of the total pulse (internal labeling +surface labeling values). All data were checked for normality with Shapiro Wilk’s test. Outliers, identified in box plots, were removed and non-normal data were natural log transformed to obtain normal distribution. Specifically four outliers were removed out of 164 data points in Fig 5d. In general, data were analysed by ANOVA, but if a co-variate was present (eg: “area” or “total pulse”) then ANCOVA was used, taking these co-variates into account.

Where applicable P-values were calculated assuming equal variances among groups (tested with Levene’s test) and were 2-sided unless stated otherwise. Differences between groups when more than two were present were identified by Bonferroni’s post-hoc test. The total number of cells per condition (n) analysed is stated on each bar and the total number of experiments given as N. All electrophysiology data were analysed with the Mann-Whitney U test using GraphPad Prism software (GraphPad) and Clampfit (Molecular Devices). Δmax describes the percentage of maximum difference between each pair of distributions. Significant statistical difference between distributions of mEPSC and mIPSC amplitudes and frequencies is defined by p< 0.01.

## Author contributions

H. B and M.E.H conceived of the project. H.B wrote the manuscript. M.E.H and C.H edited the manuscript. H.B designed and carried out all experiments and analyzed data, except the electrophysiology. P.W assisted in co-IP experiments, flow cytometry, and cloning. Z.H carried out co-IP and flow cytometry experiments and provided critical comments on the manuscript. T.F and C.H carried out slice culture electrophysiological recordings and analyzed data. M. L. provided clinical assessments of patients.

## Acknowledgments

We thank Dr Ines Ibanez-Tallon (The Rockefeller University) for the pFU-cMVIIA-PE plasmid and for helpful discussions regarding experimental design, Drs Beatriz Antolin-Fontes and Nathalie Blachere (The Rockefeller University), Kunihiro Uryu and Nadine Soplop (Rockefeller University EM Facility), Pablo Ariel, Alison North (Rockefeller University Imaging Facility), and Brian Dill (Rockefeller University Proteomics Facility) for expert advice and technical assistance, and Drs Mustafa Sahin (Boston Children’s Hospital), David Solecki (St Jude Children’s Research Hospital), and Eve Govek (The Rockefeller University) for helpful comments on the manuscript and Yin Fang for technical support. We also thank Leila Jamal and Dr Denise Batista (Kennedy Krieger Institute) for communicating details of patient CNVs and facilitating sample collection. The Neuroligin constructs were kind gifts from Drs Ann-Marie Craig (University of British Columbia) and Peter Scheiffele (University of Basel), and the SLC12a5 construct from Dr Pavel Uvarov (University of Helsinki). The Rockefeller University Proteomics Resource Center acknowledges funding from the Leona M. and Harry B. Helmsley Charitable Trust. This study makes use of data generated by the DECIPHER community. A full list of centers that contributed to the generation of the data is available from http://decipher.sanger.ac.uk and funding was provided by the Wellcome Trust.

---

**Figure S1.**
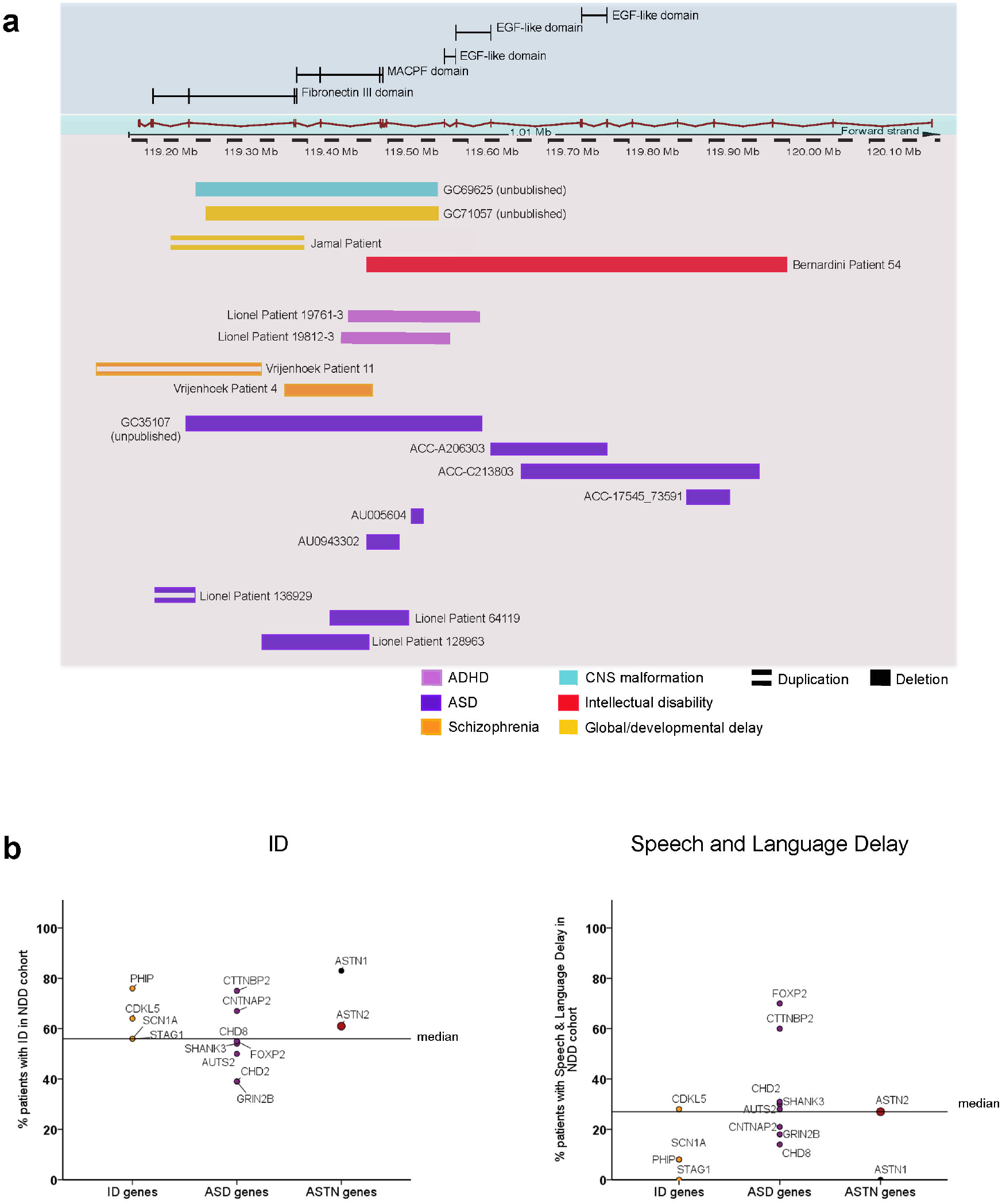
Genomic representation of ASTN2 CNVs and comparison of the occurrence of ID and speech and language impairment in patients. (**a**) Schematic of CNVs (pink box) along the *ASTN2* genomic sequence (green box) with ASTN2 protein domains encoded by each exon depicted at the top (blue box). The CNVs are color-coded according to the diagnosis of the patient (stated in the key at the bottom) with deletions represented by solid boxes and duplications by lined boxed. The CNVs represented were gathered from the following reports (Vrijenhoek et al., 2008, Glessner et al., 2009, Lionel et al., 2011, Bernardini et al., 2010) and personal communications (L. Jamal, Johns Hopkins University). For additional *ASTN2* CNVs please refer to Lionel et al., 2014 and the DECIPHER database. (**b**) Comparison of the occurrence rate of ID and speech and language delay in patients with various genetic lesions; 4 genes with strong association with ID, 8 genes with strong association with ASD, versus patients with *ASTN2* or *ASTN1* CNVs (DECIPHER database: www.decipher.sanger.ac.uk). *ASTN2* CNVs fall above the median for ID, *ASTN1* is the highest scoring for ID, while *ASTN2* but not *ASTN1* is among the highest for enrichment in speech and language delay, excluding the classical speech and language genes *FOXP2* and *CNTNAP2* (Graham and Fisher, 2013)

**Figure S2.**
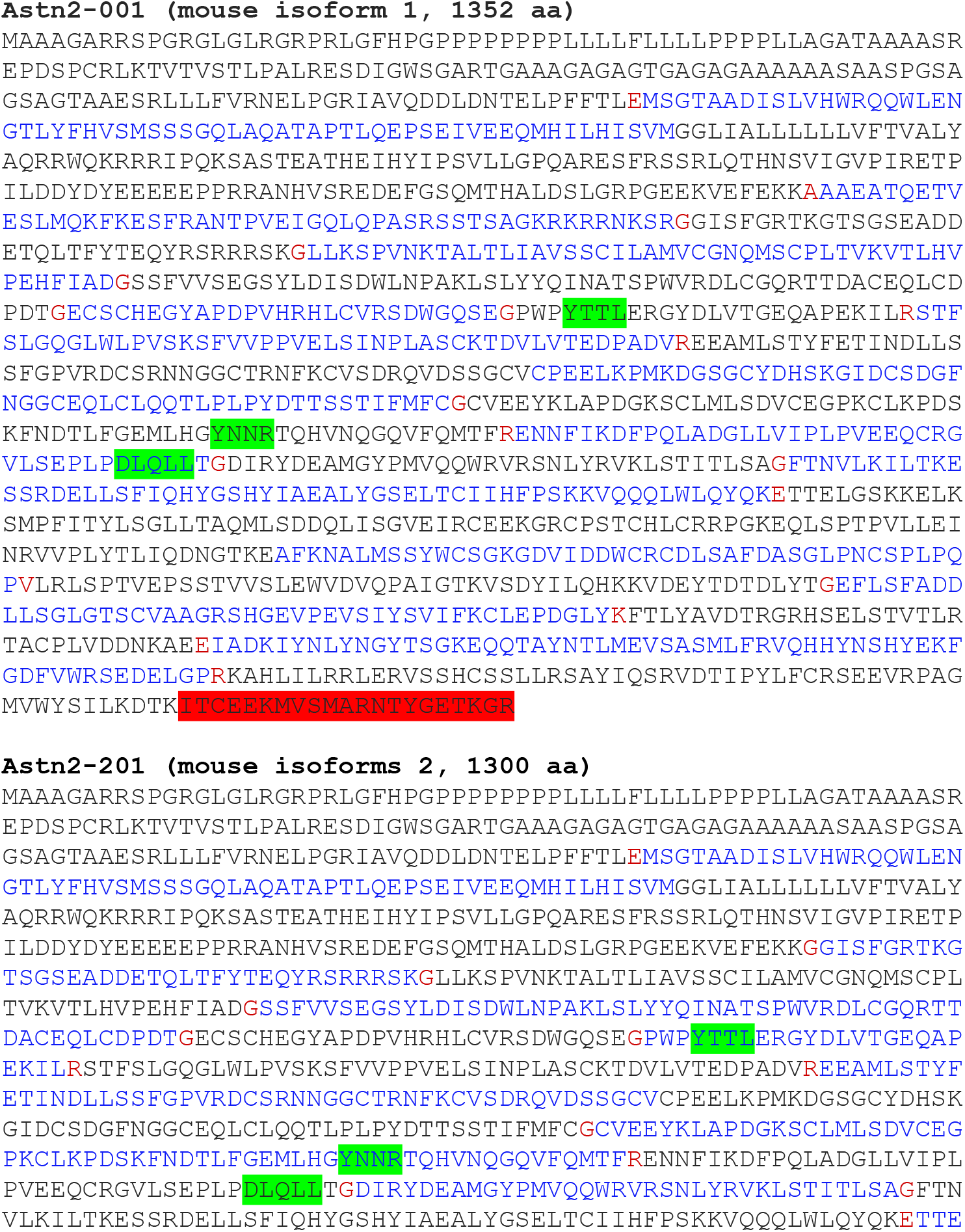
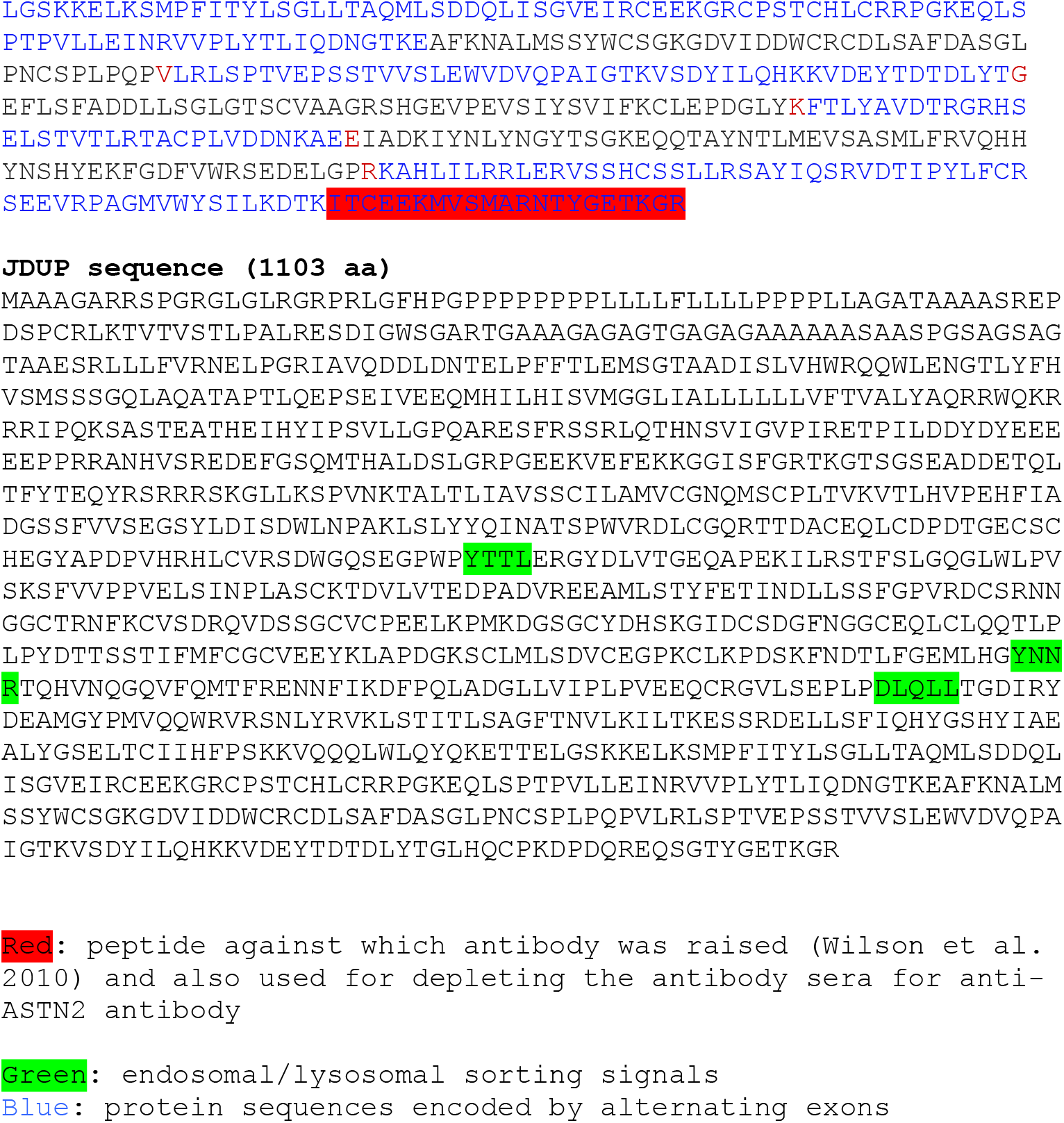
Endo/lysosomal signals in ASTN2and JDUP protein sequences. Protein sequences of the two mouse isoforms of ASTN2 and the truncated version (JDUP) modeled on the patient family CNV. Red boxes indicate the peptide against which the ASTN2 antibody was raised (Wilson et al., 2010) and also used for depleting the antibody sera for use as a control in IPs (Fig. 4 and Fig. S6). Green boxes highlight endosomal and lysosomal sorting signals. Blue texts highlight protein sequences encoded by alternating exons.

**Figure S3.**
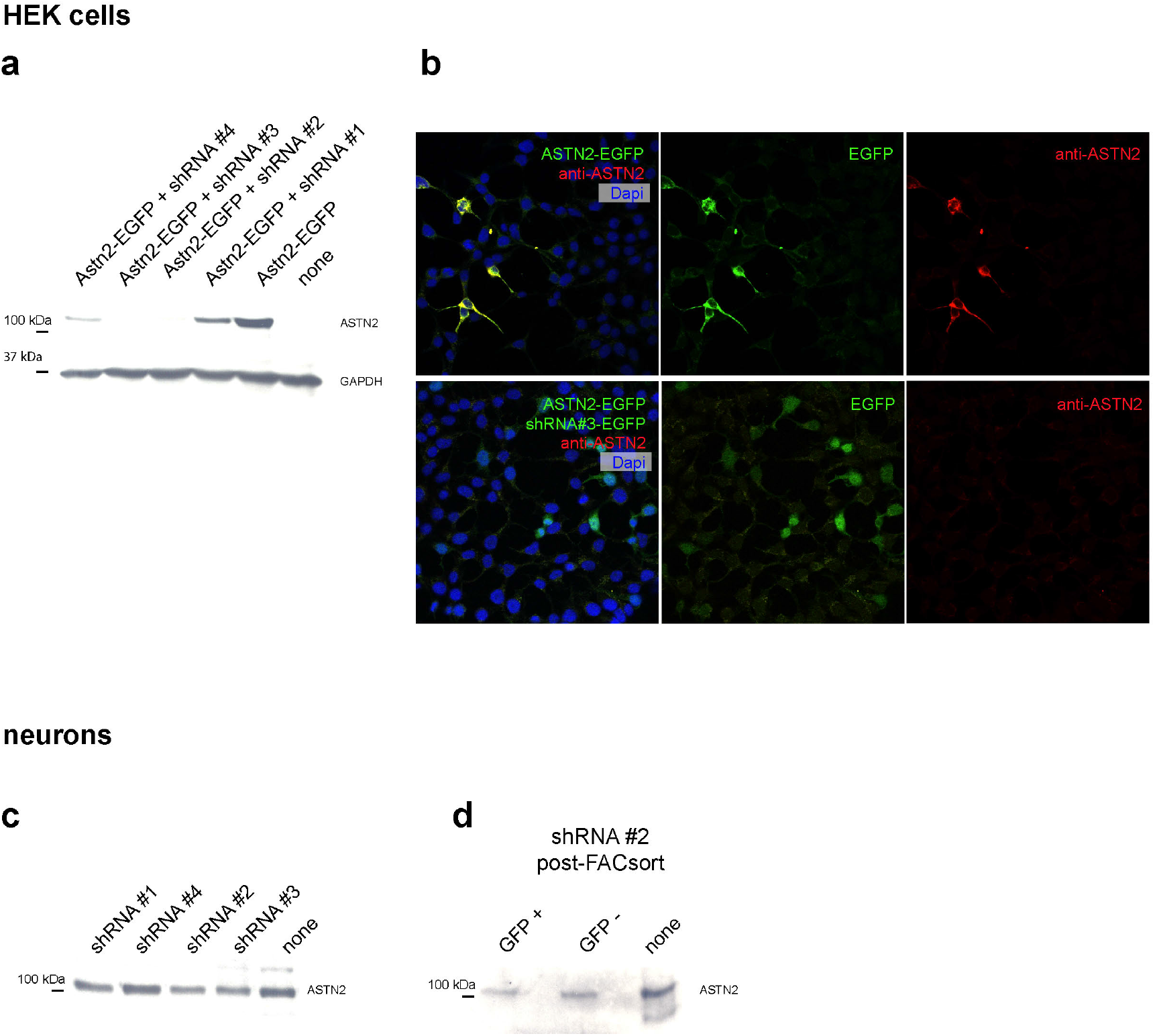
shRNA-mediated knockdown of ASTN2 and antibody specificity. (**a, b**) Knockdown of ASTN2-EGFP by four different shRNA constructs in HEK 293T cells by Western blot (a) and immunohistochemistry (b) using a rabbit antibody against ASTN2. GAPDH (at 37 kDa) was used as loading control in a. In b, top panel shows ASTN2-EGFP (green, cytoplasmic/membrane) and detected by anti-ASTN2 (red), bottom panel shows EGFP expressed from the shRNA construct (green, nuclear and cytoplasmic) but no trace of ASTN2-EGFP labeling with the ASTN2 antibody (red channel) in the presence of shRNA#3. Dapi marks nuclei. (**c, d**) Western blots showing ASTN2 protein expression in cerebellar granule cells transfected with the same shRNA constructs as in a. (c) shows Western blot on nonsorted mixed transfected cells and (d) shows lysates from FACsorted GFP-positive (expressing the shRNA construct) and GFP-negative populations.

**Figure S4.**
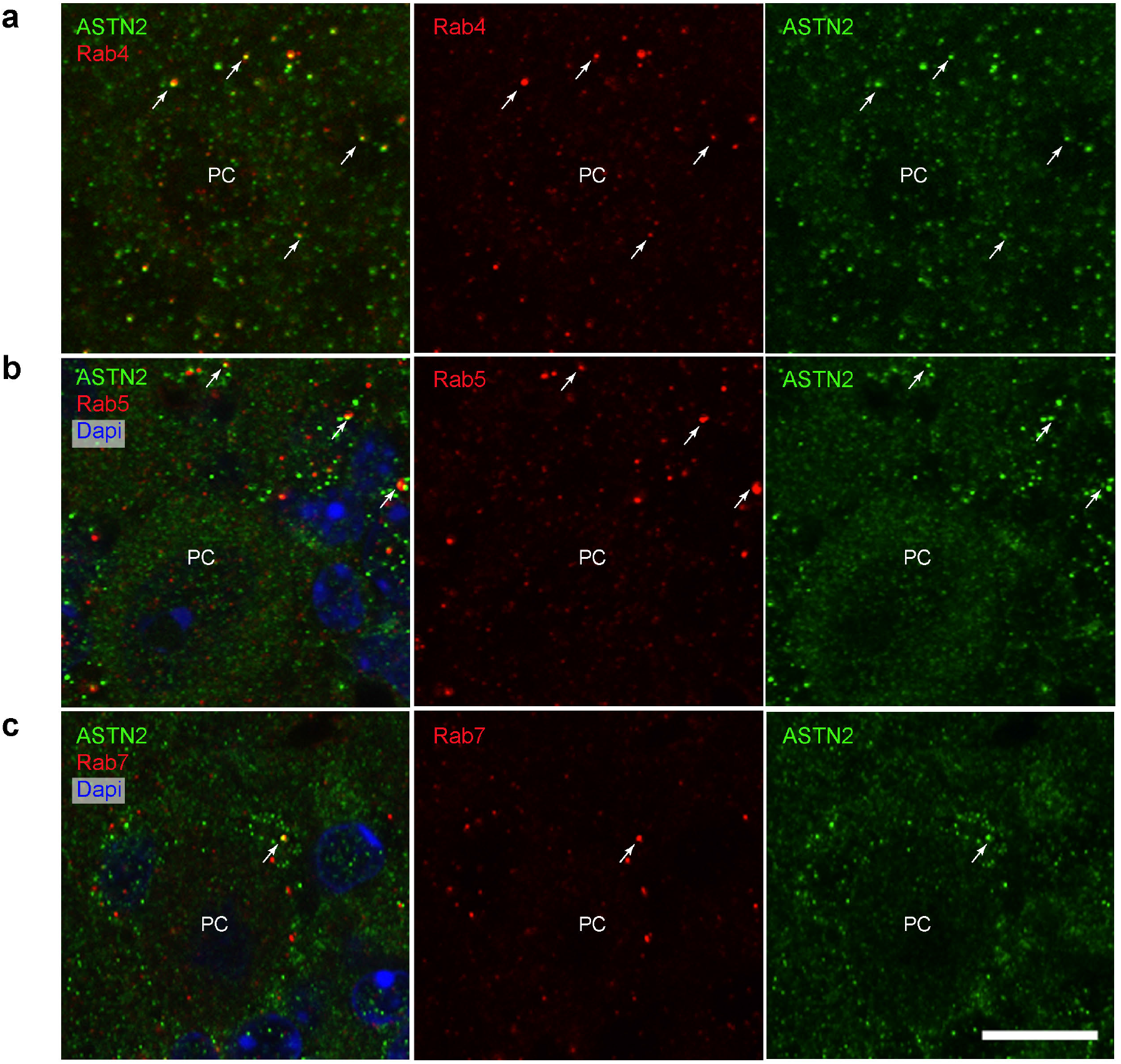
ASTN2 co-localization with endosomal markers. Immunohistochemistry of ASTN2 (green) with markers for recycling Rab4 (**a**), early Rab5 (**b**), and late Rab7 (**c**) endosomes in red in the postnatal cerebellum. Each section (sagittal) shows a PC soma and its immediate surroundings. Arrows point to examples of co-labeled puncta. Nuclei are marked by Dapi (blue) in b and c. PC, Purkinje cell, Scale bar: 10 μm

**Figure S5.**
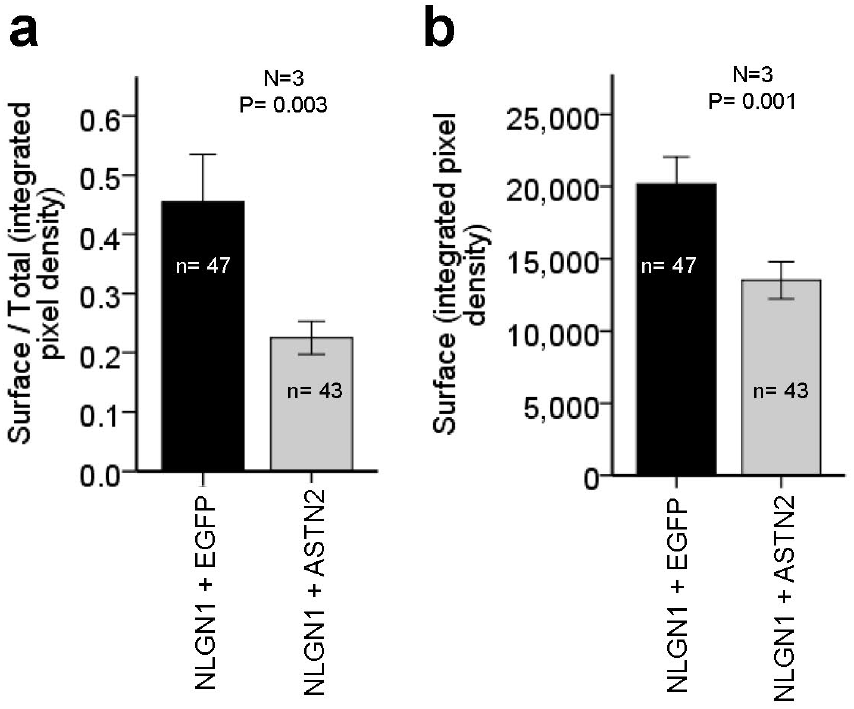
Quantification of surface labeling of NLGN1 in cerebellar granule cells. Image quantifications of surface labeling of NLGN1-HA-YFP co-expressed with either EGFP or ASTN2-EGFP in GCs. Graphs show the integrated pixel density of surface labeling as an index of total labeling (corrected for background, **a**), as well as surface labeling alone (**b**). Bars show mean +/−1 SEM from three independent experiments. The number of cells analysed per condition are stated on each bar. P-values were obtained for comparison of surface labeling by ANCOVA in a, taking into account total labeling, and by ANOVA in b.

**Figure S6.**
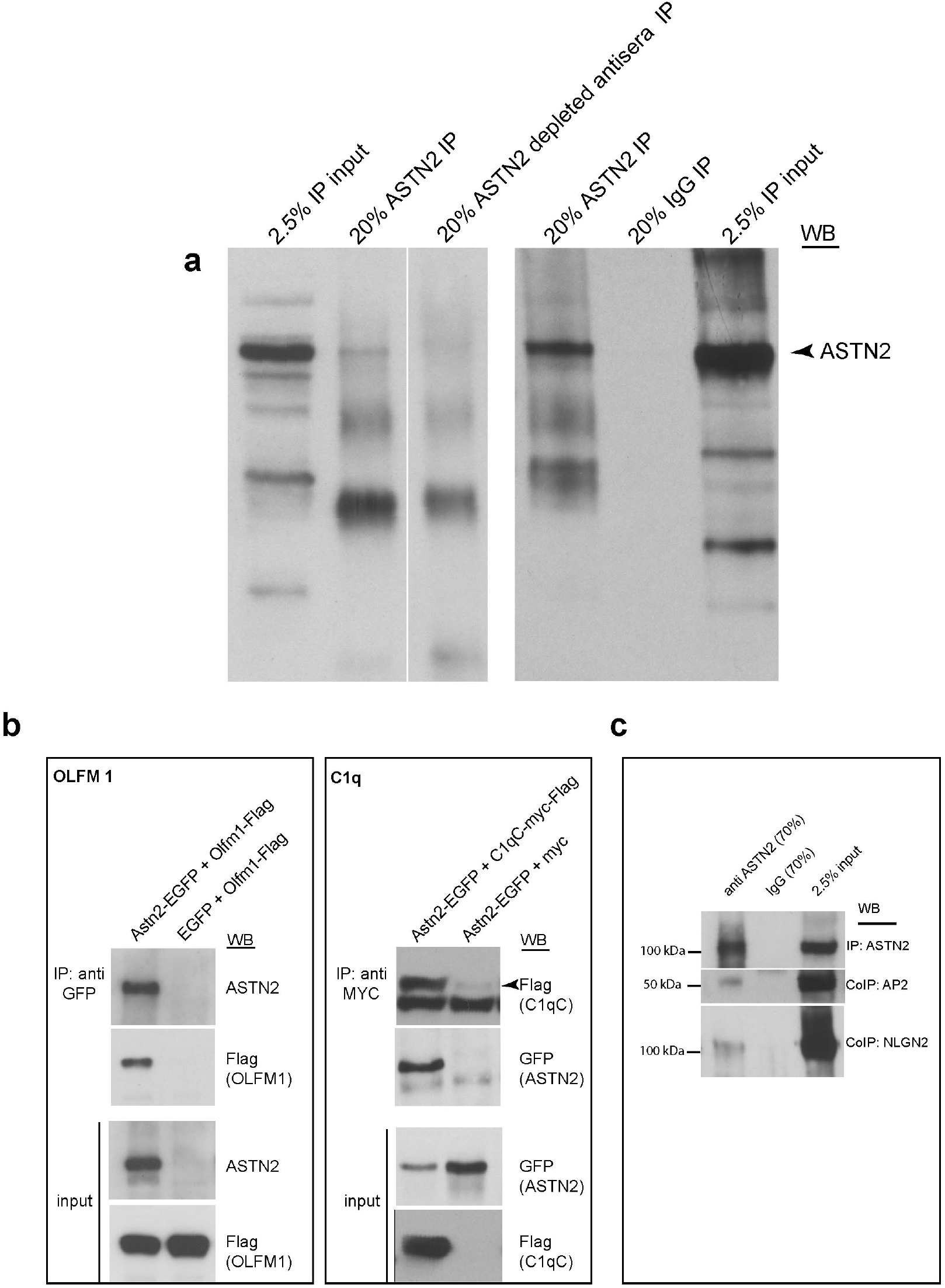
Immunoprecipitation of ASTN2 and protein interactors. (**a**) Representative Western blots of IPs with anti-ASTN2, rabbit IgG control (right panel), or depleted ASTN2 anti-sera (left panel) from the juvenile cerebellum used for mass spec analysis. Blots show 20% of total IP volumes in relation to 2.5% inputs. As the IPs processed for mass spec were directly eluted in 8M Urea from beads and were not analysed by Western blot, examples of IPs carried out with the same conditions are shown. (**b**) Co-IP of OLFM1-Flag with ASTN2-EGFP and ASTN2-EGFP with C1qc-Flag-myc in HEK293T cells. (**c**) Co-IP of AP2 and NLGN2 with ASTN2 in lysates from the juvenile cerebellum.

**Figure S7.**
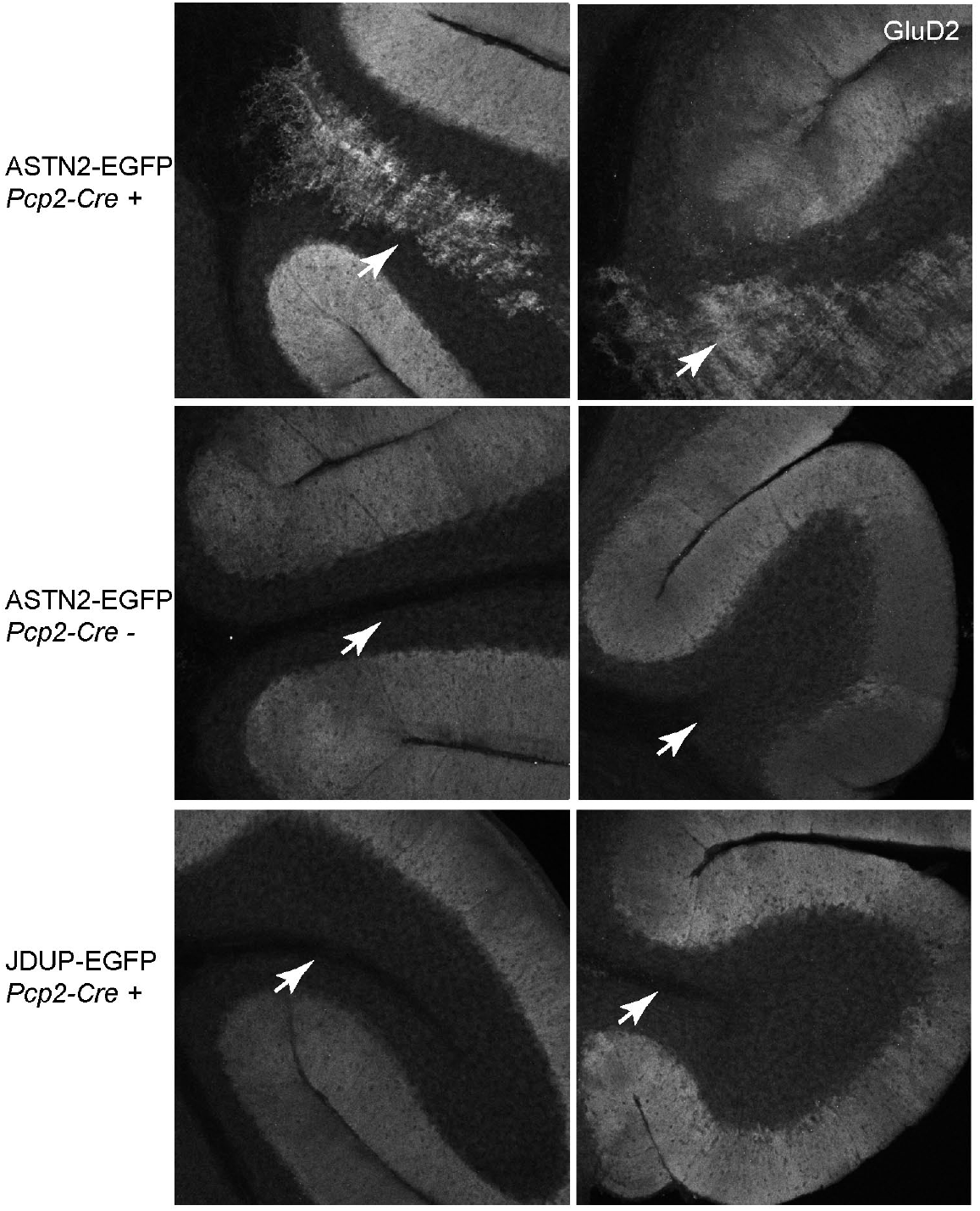
Ectopic Purkinje cells upon conditional expression of ASTN2-EGFP in the cerebellum. **(a)** Sagittal sections of the cerebellum showing ectopic PCs in the IGL and the WM (highlighted by arrows) of Lobules X (right panels) and I (left panels) marked by GluD2 expression, in *PCP2-Cre+* mice injected with pfU-cASTN2-EGFP (top), but not in pfU-cJDUP-EGFP (bottom) injected mice or in *PCP2-Cre^−/−^* mice injected with pfU-cASTN2-EGFP (middle).

## Supplementary Materials and Methods

### RT-PCR and qRT-PCR

RT-PCR was carried out according to the manufacturer’s descriptions using the HotStarTaq *PLUS* DNA Polymerase kit (Qiagen) with the following primers (*ASTN2*: forward, 5’-TACTGGTGCTCCAGGGAAAGG, reverse, 5’-CCCAATAGCTGGCTGAACAT, *β-ACTIN:* forward, 5’-AAACTGGAACGGTGAAGGTG, reverse, 5’-AGAGAAGTGGGGTGGCTTTT). qRT-PCR was performed with TaqMan primer/probe sets *(ASTN2* TaqMan gene expression assay (#Hs01024740_m1) which detects exon boundary 18-19, Human *GUSB* (Beta Glucuronidase) Endogenous Control (#4333767T, Applied Biosystems) and TaqMan Fast Advanced Master Mix, all according to the manufacturers’ descriptions on a Roche LightCycler 480 (Roche).

### Immunohisto/cytochemistry

Briefly, vibratome sections were blocked with 15% normal horse serum (Gibco), 0.1% saponin in PBS overnight and then incubated with primary antibodies overnight at 4°C and with Alexa Fluor^®^ secondary antibodies for 2 hours to overnight at room temperature and 4°C respectively. *In vitro* cultured cells were blocked in 1% normal horse serum, 0.05% Triton, incubated in primary antibodies overnight at 4°C followed by secondary Alexa Fluor^®^ antibodies for 1 hour at room temperature. Sections/cells were mounted with ProLong^®^ Gold anti-fade mounting media and sections were covered with 1.5 thickness Fisherbrand cover glass.

### Antibodies used for immunohistochemistry, immunoprecipitation and Western blot

<u>Primary antibodies, Rabbit</u>: anti-ASTN2 (1:1000-2000 on sections, and 1:500 on cells, 1:200 for Western Blot (1), anti-Calbindin D28-k (1:500, Swant #CB38), anti-GFP (1:500, Invitrogen #A11122). <u>Mouse</u>: anti-Calbindin (1:500, Swant #300), antiFlag (1:1000, Sigma #1804), anti-HA (1:500, Roche #1 583 816 001), anti-cMYC (1:50, Calbiochem #0P10), anti-AP-2 (1:250, BD Transduction Laboratories #611350), anti-NLG2 (1:100, Synaptic Systems #129 511), anti-GAPDH (1:10,000, Chemicon #mab374) anti-ROCK2 (1:1000, BD Transduction Laboratories #610623), anti-Rab4 (1:100, BD Transduction Laboratories #610888), anti-Rab5 (1:200, Synaptic Systems #108011), anti-Rab7 (1:100, Santa Cruz #sc-376362). Goat: anti-GluD2 (1:100, Santa Cruz #sc-26118). Secondary antibodies: Donkey anti-mouse, - rabbit, and -goat IgG conjugated to Alexa 405 (abcam), 555, 633, and 647 (Molecular Probes), all used at 1:300. HRP-conjugated secondary antibodies (Jackson Immunoresearch) were used at 1:8000 (anti-mouse, # 515-035-062) or 1:3000 (anti-rabbit #111-035-144) for Western blots.

### cDNA/ShRNA constructs

The following plasmids were used: pGIPZ lentiviral plasmids containing *Astn2* shRNA (# V3LMM440320) and scramble shRNA (Thermo Scientific Open Biosystems), OLFM1-MYC-Flag (Origene #MR207779), AP2s-MYC-Flag (Origene #MR200768), Clqc-MYC-Flag (Origene #MR203092), pCAG:GPI-GFP (Addgene #32601), pMES-SLC12a5-HA (50), pNice-NLGN1-CFP, pNice-NLGN2-CFP, pNice-NLGN3-YFP, pNice-NLGN4-YFP (51, 52), pNice-NLGN1-HA-YFP (gift from Dr Peter Scheiffele), pCdh2-CFP (Cdh2 cDNA gift from Dr. Richard Huganir) (53), pRK5-MYC and pMSCXβ-Venus-α-tubulin (used for flow cytometry control) were provided by Dr David Solecki. ASTN2 and JDUP containing constructs were created as follows: the sequence between the Xbal/BamHI sites of the pFU-cMVIIA-PE lentiviral plasmid (54) was removed including the DsRed/LoxP sequences. The full length *Astn2* mouse cDNA was amplified from previously reported plasmids (1) and modified to include the 5’ region of the gene, using a 5’ primer with an Xbal restriction site and a 3’ primer with a BamHI site. This sequence (full length ASTN2 splice variant 201, www.ensemble.org) was inserted in frame into the pFU backbone at Xbal/BamHI, creating pFU-Astn2-EGFP. To create the conditional construct, pFU-cAstn2-EGFP, a LoxP-dsRED-LoxP sequence (synthesized as a gblock fragment by Integrated DNA Technologies, IDT) was inserted into the Xbal site, upstream of the *Astn2* sequence. pFU-cJDUP-EGFP was created by excising the sequence between the BsiWl/BlpI sites of pFU-cAstn2-EGFP and replacing it with a synthesized sequence representing a FNIII domain deleted version (gblock, IDT). The non-conditional version (pFU-JDUP-EGFP) was created by excising the LoxP-DsRED-LoxP sequence in pFU-cJDUP-EGFP with XbaI and re-ligating the plasmid. ASTN2-HA-FLAG and JDUP-HA-FLAG plasmids were made by replacing the EGFP sequence in pFU-ASTN2-EGFP and pFU-JDUP-EGFP with in-frame HA-FLAG sequences.

### Knockdown of ASTN2 in neurons

Mixed cerebellar neurons isolated at P7 were nucleofected (Amaxa Nucleofector II) with shRNA or scrambled constructs using the mouse neuron nucleofector kit (Lonza) according to the manufacturer’s description. Cells were cultured as described previously for six days and then processed for Western blot. In a second experiment, GFP+ cells indicative of shRNA construct expression were sorted (BD FACSAria) from GFP-negative cells and then processed for Western blot.

## References

1. Wilson PM, Fryer RH, Fang Y, & Hatten ME (2010) Astn2, a novel member of the astrotactin gene family, regulates the trafficking of ASTN1 during glial-guided neuronal migration. J Neurosci 30(25):8529–8540.

2. Stitt TN & Hatten ME (1990) Antibodies that recognize astrotactin block granule neuron binding to astroglia. Neuron 5(5):639–649.

3. Adams NC, Tomoda T, Cooper M, Dietz G, & Hatten ME (2002) Mice that lack astrotactin have slowed neuronal migration. Development 129(4):965–972.

4. Fishell G & Hatten ME (1991) Astrotactin provides a receptor system for CNS neuronal migration. Development 113(3):755–765.

5. Lesch KP, et al. (2008) Molecular genetics of adult ADHD: converging evidence from genome-wide association and extended pedigree linkage studies. J Neural Transm (Vienna) 115(11):1573–1585.

6. Vrijenhoek T, et al. (2008) Recurrent CNVs disrupt three candidate genes in schizophrenia patients. Am J Hum Genet 83(4):504–510.

7. Glessner JT, et al. (2009) Autism genome-wide copy number variation reveals ubiquitin and neuronal genes. Nature 459(7246):569–573.

8. Bernardini L, et al. (2010) High-resolution SNP arrays in mental retardation diagnostics: how much do we gain? Eur J Hum Genet 18(2):178–185.

9. Lionel AC, et al. (2011) Rare copy number variation discovery and crossdisorder comparisons identify risk genes for ADHD. Sci Transl Med 3(95):95ra75.

10. Lionel AC, et al. (2014) Disruption of the ASTN2/TRIM32 locus at 9q33.1 is a risk factor in males for autism spectrum disorders, ADHD and other neurodevelopmental phenotypes. Hum Mol Genet 23(10):2752–2768.

11. Glickstein M (2007) What does the cerebellum really do? Curr Biol 17(19):R824–827.

12. Timmann D & Daum I (2007) Cerebellar contributions to cognitive functions: a progress report after two decades of research. Cerebellum 6(3): 159–162.

13. Strick PL, Dum RP, & Fiez JA (2009) Cerebellum and nonmotor function. Annu Rev Neurosci 32:413–434.

14. Fatemi SH, et al. (2012) Consensus paper: pathological role of the cerebellum in autism. Cerebellum 11(3):777–807.

15. Kloth AD, et al. (2015) Cerebellar associative sensory learning defects in five mouse autism models. Elife 4:e06085.

16. Tsai PT, et al. (2012) Autistic-like behaviour and cerebellar dysfunction in Purkinje cell Tsc1 mutant mice. Nature 488(7413):647–651.

17. Price JC, Guan S, Burlingame A, Prusiner SB, & Ghaemmaghami S (2010) Analysis of proteome dynamics in the mouse brain. Proc Natl Acad Sci U S A 107(32):14508–14513.

18. Stevens B, et al. (2007) The classical complement cascade mediates CNS synapse elimination. Cell 131(6):1164–1178.

19. Schwenk J, et al. (2012) High-resolution proteomics unravel architecture and molecular diversity of native AMPA receptor complexes. Neuron 74(4):621–633.

20. Krishnan A, et al. (2016) Genome-wide prediction and functional characterization of the genetic basis of autism spectrum disorder. Nat Neurosci 19:1454–1462.

21. Zhou Z, Meng Y, Asrar S, Todorovski Z, & Jia Z (2009) A critical role of Rho-kinase ROCK2 in the regulation of spine and synaptic function. Neuropharmacology 56(1):81–89.

22. Rivera C, et al. (1999) The K+/Cl-co-transporter KCC2 renders GABA hyperpolarizing during neuronal maturation. Nature 397(6716):251–255.

23. Li H, et al. (2007) KCC2 interacts with the dendritic cytoskeleton to promote spine development. Neuron 56(6):1019–1033.

24. Merner ND, et al. (2015) Regulatory domain or CpG site variation in SLC12A5, encoding the chloride transporter KCC2, in human autism and schizophrenia. Front Cell Neurosci 9:386.

25. Tang X, et al. (2016) KCC2 rescues functional deficits in human neurons derived from patients with Rett syndrome. Proc Natl Acad Sci U S A 113(3):751–756.

26. Banerjee A, et al. (2016) Jointly reduced inhibition and excitation underlies circuit-wide changes in cortical processing in Rett syndrome. Proc Natl Acad Sci US A 113(46):E7287–E7296.

27. Sudhof TC (2008) Neuroligins and neurexins link synaptic function to cognitive disease. Nature 455(7215):903–911.

28. Prelich G (2012) Gene overexpression: uses, mechanisms, and interpretation. Genetics 190(3):841–854.

29. Zhang B, et al. (2015) Neuroligins Sculpt Cerebellar Purkinje-Cell Circuits by Differential Control of Distinct Classes of Synapses. Neuron 87(4):781–796.

30. Heo S, et al. (2018) Identification of long-lived synaptic proteins by proteomic analysis of synaptosome protein turnover. Proc Natl Acad Sci U S A 115(16):E3827–E3836.

31. Ni T, Harlos K, & Gilbert R (2016) Structure of astrotactin-2: a conserved vertebrate-specific and perforin-like membrane protein involved in neuronal development. Open Biol 6(5):160053.

32. Bonifacino JS & Traub LM (2003) Signals for sorting of transmembrane proteins to endosomes and lysosomes. Annu Rev Biochem 72:395–447.

33. Otomo A, Pan L, & Hadano S (2012) Dysregulation of the autophagy-endolysosomal system in amyotrophic lateral sclerosis and related motor neuron diseases. Neurol Res Int 2012:498–428.

34. Razi M, Chan EY, & Tooze SA (2009) Early endosomes and endosomal coatomer are required for autophagy. J Cell Biol 185(2):305–321.

35. Bayes A, et al. (2012) Comparative study of human and mouse postsynaptic proteomes finds high compositional conservation and abundance differences for key synaptic proteins. PLoS One 7(10):e46683.

36. Walmsley B, Alvarez FJ, & Fyffe RE (1998) Diversity of structure and function at mammalian central synapses. Trends Neurosci 21(2):81–88.

37. Khalilov I, et al. (2011) Enhanced Synaptic Activity and Epileptiform Events in the Embryonic KCC2 Deficient Hippocampus. Front Cell Neurosci 5:23.

38. Marrocco J, et al. (2017) A sexually dimorphic pre-stressed translational signature in CA3 pyramidal neurons of BDNF Val66Met mice. Nat Commun 8(1):808.

39. Allen G & Courchesne E (2003) Differential effects of developmental cerebellar abnormality on cognitive and motor functions in the cerebellum: an fMRI study of autism. Am J Psychiatry 160(2):262–273.

40. Amaral DG, Schumann CM, & Nordahl CW (2008) Neuroanatomy of autism. Trends Neurosci 31(3):137–145.

41. Stoodley CJ, et al. (2017) Altered cerebellar connectivity in autism and cerebellar-mediated rescue of autism-related behaviors in mice. Nat Neurosci 20(12):1744–1751.

42. Willsey AJ, et al. (2013) Coexpression networks implicate human midfetal deep cortical projection neurons in the pathogenesis of autism. Cell 155(5):997–1007.

43. Parikshak NN, et al. (2013) Integrative functional genomic analyses implicate specific molecular pathways and circuits in autism. Cell 155(5):1008–1021.

44. Koekkoek SK, et al. (2005) Deletion of FMR1 in Purkinje cells enhances parallel fiber LTD, enlarges spines, and attenuates cerebellar eyelid conditioning in Fragile X syndrome. Neuron 47(3):339–352.

45. Baudouin SJ, et al. (2012) Shared synaptic pathophysiology in syndromic and nonsyndromic rodent models of autism. Science 338(6103):128–132.

46. Reith RM, et al. (2013) Loss of Tsc2 in Purkinje cells is associated with autistic-like behavior in a mouse model of tuberous sclerosis complex. Neurobiol Dis 51:93–103.

47. Piochon C, et al. (2014) Cerebellar plasticity and motor learning deficits in a copy-number variation mouse model of autism. Nat Commun 5:5586.

48. Sztainberg Y & Zoghbi HY (2016) Lessons learned from studying syndromic autism spectrum disorders. Nat Neurosci 19(11):1408–1417.

49. Baptista CA, Hatten ME, Blazeski R, & Mason CA (1994) Cell-cell interactions influence survival and differentiation of purified Purkinje cells in vitro. Neuron 12(2):243–260.

50. Acton BA, et al. (2012) Hyperpolarizing GABAergic transmission requires the KCC2 C-terminal ISO domain. J Neurosci 32(25):8746–8751.

51. Graf ER, Zhang X, Jin SX, Linhoff MW, & Craig AM (2004) Neurexins induce differentiation of GABA and glutamate postsynaptic specializations via neuroligins. Cell 119(7):1013–1026.

52. Pettem KL, Yokomaku D, Takahashi H, Ge Y, & Craig AM (2013) Interaction between autism-linked MDGAs and neuroligins suppresses inhibitory synapse development. J Cell Biol 200(3):321–336.

53. Nuriya M & Huganir RL (2006) Regulation of AMPA receptor trafficking by N-cadherin. JNeurochem 97(3):652–661.

54. Auer S, et al. (2010) Silencing neurotransmission with membrane-tethered toxins. Nat Methods 7(3):229–236.

55. Kall L, Canterbury JD, Weston J, Noble WS, & MacCoss MJ (2007) Semisupervised learning for peptide identification from shotgun proteomics datasets. Nat Methods 4(11):923–925.

56. Kim JY, et al. (2013) Viral transduction of the neonatal brain delivers controllable genetic mosaicism for visualising and manipulating neuronal circuits in vivo. Eur J Neurosci 37(8):1203–1220.

57. Dugue GP, Dumoulin A, Triller A, & Dieudonne S (2005) Target-dependent use of co-released inhibitory transmitters at central synapses. J Neurosci 25(28):6490–6498.

## References

Bernardini, L., Alesi, V., Loddo, S., Novelli, A., Bottillo, I., Battaglia, A., Digilio, M. C., Zampino, G., Ertel, A., Fortina, P., Surrey, S. & Dallapiccola, B. (2010) High-resolution SNP arrays in mental retardation diagnostics: how much do we gain? EurJHum Genet, 18, 178–85.

Glessner, J. T., Wang, K., Cai, G., Korvatska, O., Kim, C. E., Wood, S., Zhang, H., Estes, A., Brune, C. W., Bradfield, J. P., Imielinski, M., Frackelton, E. C., Reichert, J., Crawford, E. L., Munson, J., Sleiman, P. M., Chiavacci, R., Annaiah, K., Thomas, K., Hou, C., Glaberson, W., Flory, J., Otieno, F., Garris, M., Soorya, L., Klei, L., Piven, J., Meyer, K. J., Anagnostou, E., Sakurai, T., Game, R. M., Rudd, D. S., Zurawiecki, D., Mcdougle, C. J., Davis, L. K., Miller, J., Posey, D. J., Michaels, S., Kolevzon, A., Silverman, J. M., Bernier, R., Levy, S. E., Schultz, R. T., Dawson, G., Owley, T., Mcmahon, W. M., Wassink, T. H., Sweeney, J. A., Nurnberger, J. I., Coon, H., Sutcliffe, J. S., Minshew, N. J., Grant, S. F., Bucan, M., Cook, E. H., Buxbaum, J. D., Devlin, B., Schellenberg, G. D. & Hakonarson, H. (2009) Autism genome-wide copy number variation reveals ubiquitin and neuronal genes. Nature, 459, 569–73.

Graham, S. A. & Fisher, S. E. (2013) Decoding the genetics of speech and language. Curr Opin Neurobiol, 23, 43–51.

Lionel, A. C., Crosbie, J., Barbosa, N., Goodale, T., Thiruvahindrapuram, B., Rickaby, J., Gazzellone, M., Carson, A. R., Howe, J. L., Wang, Z., Wei, J., Stewart, A. F., Roberts, R., Mcpherson, R., Fiebig, A., Franke, A., Schreiber, S., Zwaigenbaum, L., Fernandez, B. A., Roberts, W., Arnold, P. D., Szatmari, P., Marshall, C. R., Schachar, R. & Scherer, S. W. (2011) Rare copy number variation discovery and cross-disorder comparisons identify risk genes for ADHD. Sci Transl Med, 3, 95ra75.

Vrijenhoek, T., Buizer-Voskamp, J. E., Van Der Stelt, I., Strengman, E., Sabatti, C., Geurts Van Kessel, A., Brunner, H. G., Ophoff, R. A. & Veltman, J. A. (2008) Recurrent CNVs disrupt three candidate genes in schizophrenia patients. Am J Hum Genet, 83, 504–10.

Wilson, P. M., Fryer, R. H., Fang, Y. & Hatten, M. E. (2010) Astn2, a novel member of the astrotactin gene family, regulates the trafficking of ASTN1 during glial-guided neuronal migration. J Neurosci, 30, 8529–40.

